# RiboDFM: Robust RNA sequence design using backbone-conditioned discrete flow matching

**DOI:** 10.64898/2026.10.22.761295

**Authors:** Xiaocheng Liu, Xiaoyu An, Zhenling Peng, Wenkai Wang, Jianyi Yang

## Abstract

Designing RNA sequences that fold into predefined 3D structures is central to RNA engineering, yet it remains challenging due to the scarcity of experimentally determined structures and the complex sequence-structure relationships. Here, we present RiboDFM, a backbone-conditioned discrete flow-matching framework for RNA sequence design. RiboDFM progressively resolves fully masked sequences by integrating a geometric backbone encoder, an auxiliary base-pairing prediction head, and language model representations into a discrete flow sampler. To expand our training data, we supplemented experimental structures with a large-scale collection of trRosettaRNA2-predicted models. Across diverse benchmarks, RiboDFM achieves high sequence recovery and generates sequences with greater tertiary-structure self-consistency than existing methods. RiboDFM also performs strongly on aptamer-focused design tasks. Finally, beyond sequence generation, the nucleotide probabilities derived from RiboDFM’s flow trajectory serve as an intrinsic measure of sequence-structure compatibility. This enables reliable confidence estimation for designed sequences, as well as reference-free, zero-shot assessment of predicted structure models. Together, these results establish RiboDFM as an effective framework for structure-based RNA sequence design.

## Introduction

RNA sequence design seeks to solve the inverse folding problem: identifying nucleotide sequences that reliably fold into a predefined target conformation. Reliable RNA design therefore provides a foundation for engineering functional RNAs for molecular sensing, programmable gene regulation, and therapeutic applications[1–3]. Most conventional approaches formulate this problem at the secondary-structure level[4–6]. However, secondary structure alone does not fully specify the noncanonical base pairs, tertiary contacts, junction geometries, and global spatial organization that stabilize complex RNA folds. This limitation has motivated the development of sequence-design methods conditioned on 3D RNA backbones.

Recent advances in geometric deep learning have enabled RNA sequence design directly from 3D structures. RhoDesign employs geometric vector perceptrons (GVPs) to encode structural geometry for sequence generation [7]. gRNAde adopts a coarse-grained three-bead representation of structures and uses a multistate geometric graph neural network to design sequences compatible with either a single structure or multiple conformational states[8]. More recently, NA-MPNN extended message-passing architectures originally developed for protein design[9, 10] to nucleic acids using a more detailed backbone representation[11]. AlignIF augments the target structure by integrating structural information across similar folds[12]. Together, these approaches have advanced 3D structure-based sequence design. Nevertheless, 3D structure-based RNA sequence design is still in its infancy and far from satisfactory.

A major limitation is the scarcity of experimentally determined RNA structures relative to the rapidly expanding number of known RNA sequences. Existing structural datasets cover only a small and unevenly distributed fraction of RNA sequence and fold space, restricting the structural diversity available for supervised learning[13–17]. Recent improvements in RNA structure prediction offer a potential route to alleviating this limitation[18–26]. Methods such as trRosettaRNA[18] and trRosettaRNA2[20] can generate 3D structural models for large collections of RNA sequences that lack experimentally resolved structures, thereby providing an additional source of sequence-structure pairs for model training. Although predicted structures inevitably contain modeling errors, carefully curated predicted models may broaden the range of folds and sequence contexts encountered during training.

Another challenge concerns the sequence-generation framework. Most existing 3D RNA design methods use autoregressive decoding, factorizing the conditional sequence distribution into a series of single-position predictions according to a predefined or randomly sampled order[7, 8, 11]. Although highly effective, this formulation requires sequential decoding over a number of steps that scales with RNA length, which can increase computational cost for longer RNAs. It also imposes a particular factorization of the sequence distribution, with each prediction conditioned on nucleotide choices made at earlier positions. Such sequential dependence can be restrictive for RNA, in which base pairing, tertiary contacts, and compensatory substitutions couple residues that may be widely separated along the sequence. A generative framework that can simultaneously update multiple positions is required.

To address the above challenges, we developed RiboDFM, a backbone-conditioned discrete flow-matching framework [27] for RNA sequence design. Unlike strictly position-by-position autoregressive decoding, RiboDFM predicts nucleotide distributions for all positions at each flow time. Comprehensive evaluations across complementary benchmarks show that RiboDFM outperforms existing methods in both sequence recovery and structure consistency.

## Results

### Overview of RiboDFM

As shown in Fig. 1a, RiboDFM is a discrete flow-matching framework [27] for RNA sequence design conditioned on a target backbone structure. Given a target RNA backbone with unspecified base identities, RiboDFM first constructs a nucleotide-level *k*-nearest-neighbor (*k*-NN) graph using the C4′ atoms as residue centers. Graph edges are encoded with geometric features, including inter-nucleotide distances and relative positional information along the sequence (see Methods). The resulting graph is then processed by a message-passing neural network (MPNN) [28] encoder to produce node- and edge-level representations of the target backbone. Base-pair probabilities are further inferred from the learned edge representations by a secondary-structure (SS) head. Finally, the backbone representations and predicted base-pair probabilities condition the subsequent discrete flow-matching sampling process, which progressively generates an RNA sequence compatible with the target backbone.

**Figure 1.**
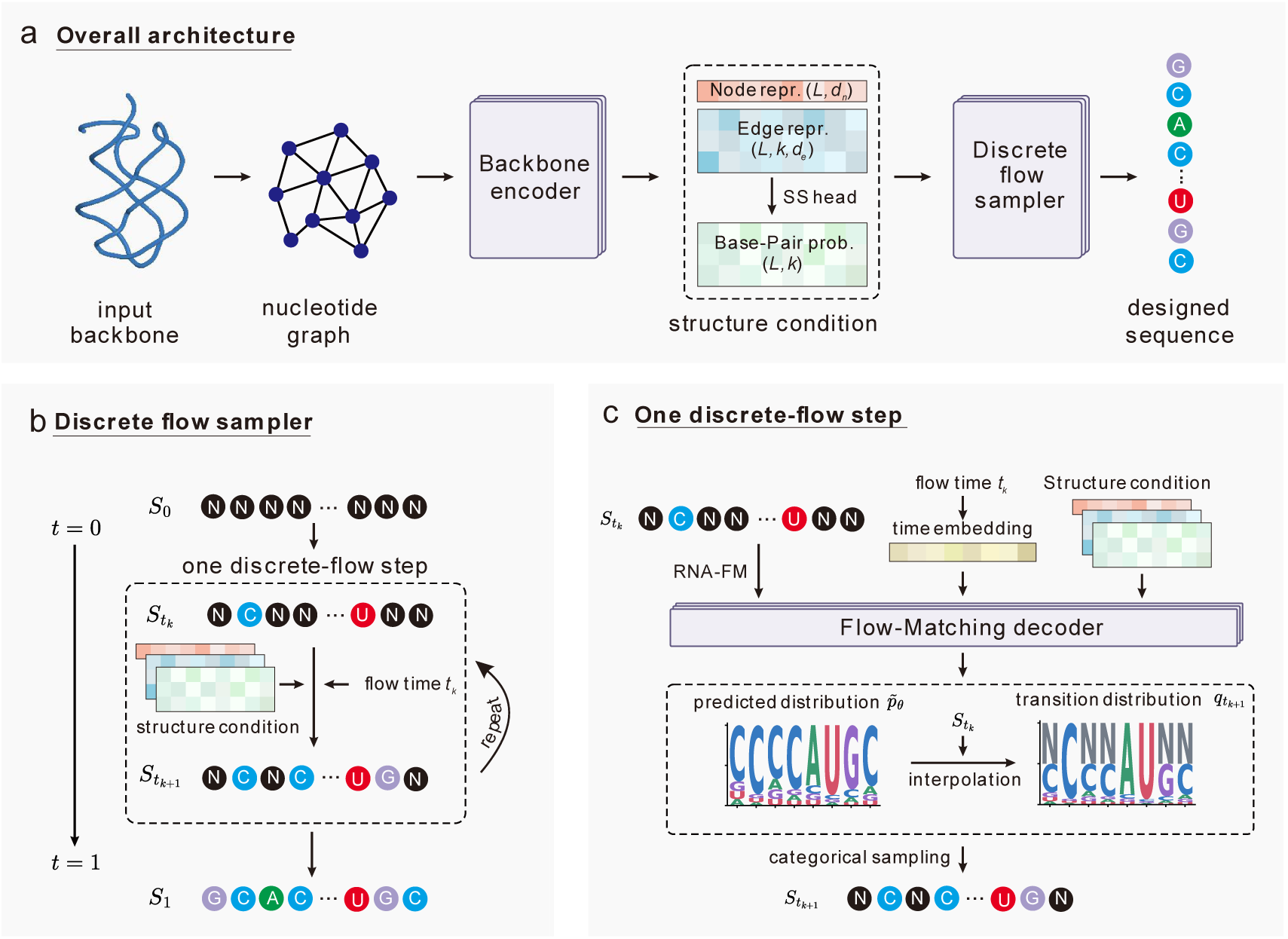
Overview of the RiboDFM framework for backbone-conditioned RNA sequence design. **a.** Overall architecture of RiboDFM. The input RNA backbone is represented as a nucleotide-level graph and processed by the backbone encoder to produce node and edge representations. An auxiliary secondary-structure (SS) head predicts base-pairing probabilities from the edge representations. These features provide the structural condition that guides the discrete flow sampler in generating an RNA sequence. *L* denotes the sequence length, *k*=128 specifies the maximum number of nearest neighbors per nucleotide, and *d_n_* and *d_e_* denote the node and edge feature dimensions, respectively, both set to 256. **b.** Discrete flow sampling trajectory. Starting from the fully masked sequence *S*_0_, RiboDFM iteratively updates and refines the sequence state over a predefined number of flow steps (32 by default). The state obtained after the final step at *t*=1 is taken as the designed sequence *S*_1_, with multiple positions processed in parallel at each step. **c.** A single flow step. At time *t_k_*, the flow-matching decoder integrates the RNA-FM representation of the current sequence state *S_tk_* , the structural condition, and the flow-time embedding to predict position-wise nucleotide distributions *p_θ_* . These predicted distributions are interpolated with the current sequence state to construct the transition distributions *q_tk_* , from which the next sequence state *S_tk_* is sampled.

The core of RiboDFM is an RNA sequence sampler based on backbone-conditioned discrete flow matching (Fig. 1b). Conditioned on the target backbone, the discrete flow sampler progressively transforms a fully masked sequence into a complete RNA sequence. Specifically, at each flow step (Fig. 1c), the current sequence state is first encoded by the pretrained RNA language model RNA-FM[29]. The resulting RNA-FM embedding, together with a Fourier embedding of the flow time and the backbone representations from the encoder, is then fed into the flow-matching decoder, which consists of three MPNN blocks. The decoder predicts a probability distribution over the four nucleotide types (A, U, C, and G) for each residue. These predicted distributions are then interpolated with the current sequence state using a time-dependent weight determined by the flow time *t*, yielding position-specific transition distributions. Each position is independently sampled from its corresponding transition distribution, allowing the current nucleotide state to be either retained or updated. More details about model architecture and training implementation are provided in Methods.

### RiboDFM improves RNA sequence design across diverse benchmarks

To systematically evaluate RiboDFM, we benchmarked it on three complementary RNA sequence design datasets: Mono63, CASP16, and New38 (see Methods for details). Mono63, originally curated by NA-MPNN, comprises 63 monomeric RNAs and represents a standard benchmark for backbone-conditioned RNA design, whereas CASP16 contains 25 challenging targets from the community-wide blind CASP16 structure-prediction experiment[30]. Both datasets were filtered to exclude targets with sequences redundant to those in our training set. To further assess generalization to recently released RNAs while minimizing potential overlap with the training data of baseline methods, we constructed New38, an additional benchmark comprising 38 RNA structures released in the PDB after 21 January 2025.

Beyond sequence recovery (the sequence identity between the designed and native sequences), we evaluated tertiary-structural self-consistency to provide a more comprehensive assessment of design quality. Specifically, for each designed sequence, we predicted its 3D structures and compared them with the corresponding target structures. To reduce dependence on any single predictor and improve the robustness of the structural self-consistency evaluation, we independently predicted the 3D structure of each designed sequence using three methods: trRosettaRNA2[20], AlphaFold 3[31], and Protenix[32]. Among all predicted structures, the one with the lowest C1’-RMSD to the target backbone was selected as the representative prediction for the designed sequence. All subsequent 3D self-consistency metrics, including lDDT, TM-score, RMSD, and INF, were then calculated using this same selected structure and are denoted as sclDDT, scTM, scRMSD, and scINF, respectively (following the self-consistency “sc” notation used in previous RNA design work[8]).

We benchmarked RiboDFM against three representative methods: NA-MPNN[11], gRNAde[8], and RhoDesign[7]. Because nucleobase identities are inherently unknown during RNA sequence design, utilizing any base-derived atomic coordinates risks information leakage. Both RiboDFM and NA-MPNN rely strictly on sugar-phosphate backbone coordinates. Conversely, RhoDesign additionally incorporates base-specific N1 and C2 atoms. This inclusion may risk information leakage during sequence design by inadvertently providing the model with binary purine/pyrimidine hints. Although gRNAde treats N1/N9 coordinates in a base-agnostic manner to avoid this direct leakage, utilizing these atoms may still inadvertently leak structural information about the nucleobases.

We followed the evaluation protocol used in NA-MPNN. For each target, each method was run independently with ten different random seeds, generating one sequence per run at a sampling temperature of 0.1. Performance was summarized under two complementary settings: (1) Average, in which the target-level score was calculated as the mean metric value across the ten runs, reflecting the method’s expected performance under repeated sampling; and (2) Top1, in which the sequence-structure pair with the lowest scRMSD among the ten runs was selected for each target, and all metrics, including sequence recovery, sclDDT, scTM, scRMSD, and scINF, were reported for that pair, reflecting the quality of the most structurally consistent candidate generated within a fixed sampling budget. To avoid redundancy, subsequent discussion focuses primarily on the Average results unless otherwise noted, as the two evaluation settings yield broadly consistent comparative trends.

As shown in Fig. 2a and Supplementary Table 1, RiboDFM consistently achieves strong performance across all the evaluated benchmarks, particularly in terms of 3D structural self-consistency. On the particularly challenging CASP16 benchmark, for example, RiboDFM improves the average sclDDT from 46.6 to 49.3 and scRMSD from 28.3 to 27.5 Å relative to the recent state-of-the-art method NA-MPNN. Target-wise head-to-head comparisons further show that these gains are broadly distributed across all three benchmark sets rather than being driven by a small subset of targets (Fig. 2b). Notably, the advantage is especially pronounced on New38, which was constructed from recently released RNA structures to minimize potential overlap with the training data of baseline methods, providing further evidence that RiboDFM generalizes well to newly released RNAs.

**Figure 2.**
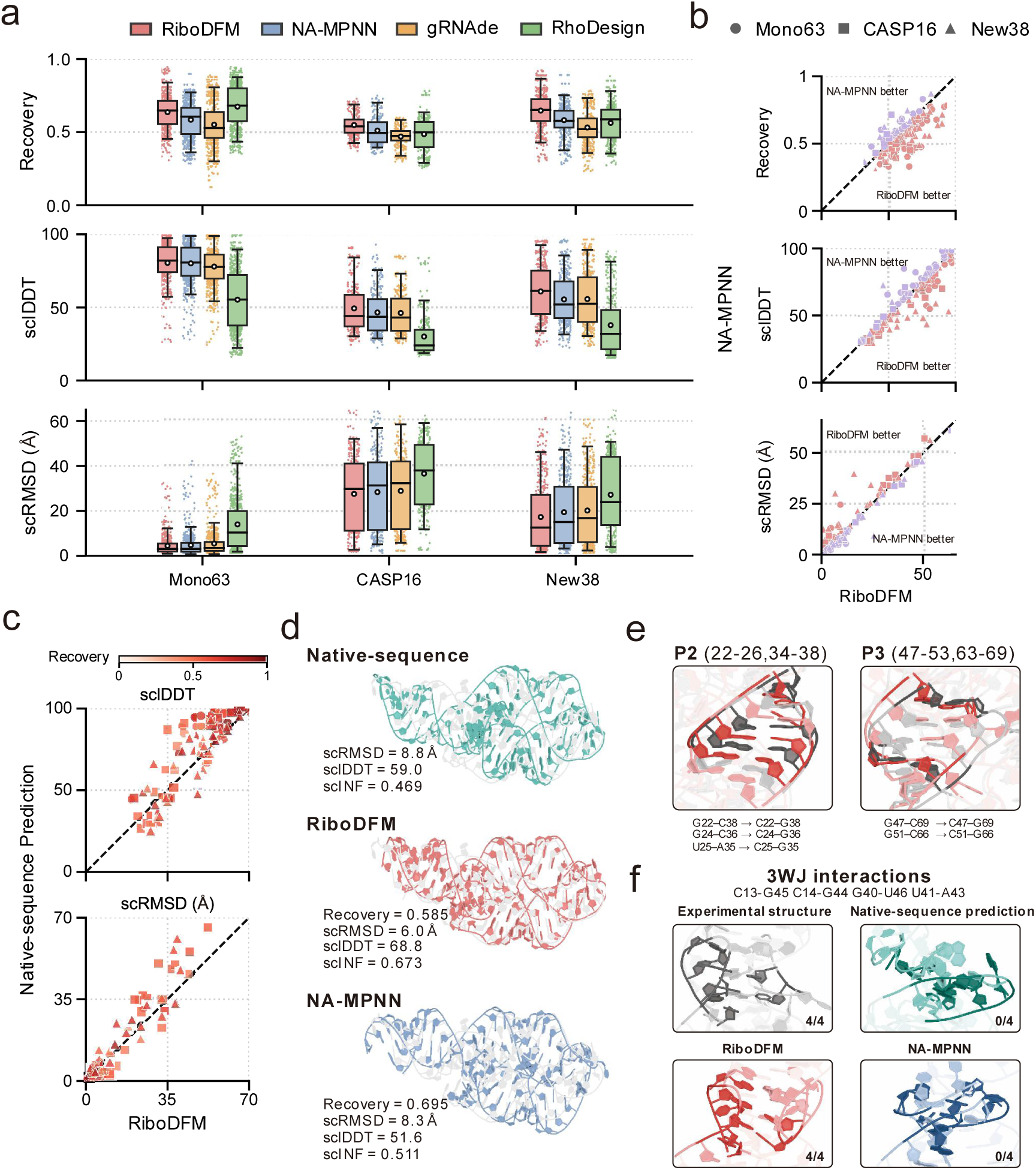
Performance of RiboDFM compared with other RNA design methods. **a**, Distributions of sequence recovery, sclDDT and scRMSD across ten designed sequences per target on the Mono63, CASP16 and New38 benchmarks. **b.** Target-wise comparisons between RiboDFM and NA-MPNN under the Average setting, where each metric was averaged across the ten designs generated for each target. **c.** Comparison of the scRMSD-top1 RiboDFM designs with predictions obtained from the corresponding native sequences. Colors indicate the sequence recovery of the RiboDFM designs. **d-f,** Representative example for target 12CI, using the scRMSD-top1 design for each method. **d,** Structural comparison of the predicted structures from the native sequence and from sequences designed by RiboDFM and NA-MPNN; **e,** Compensatory base-pair substitutions in P2 and P3 stems of the RiboDFM design. **f,** Recovery of four selected native interactions supporting the three-way junction (3WJ) in the experimental structure and in predictions from the native sequence, RiboDFM design, and NA-MPNN design.

In terms of sequence recovery, RiboDFM also achieves the best performance on CASP16 and New38, although it performs slightly worse than RhoDesign on Mono63 (0.637 vs. 0.675). However, RhoDesign’s higher recovery does not translate into greater structural fidelity, as it performs substantially worse on 3D structure self-consistency metrics. This discrepancy can be attributed to two factors. First, Mono63 was filtered to remove sequence redundancy with the training sets of RiboDFM and NA-MPNN, but not with those of gRNAde and RhoDesign; therefore, potential overlap with the training data of the latter two methods was not ruled out. Second, RhoDesign appears effective at distinguishing purines from pyrimidines but frequently confuses nucleotides within the same class (Supplementary Fig. 1). In particular, it misclassifies 41.0% of native G nucleotides as A, compared with only 9.9%, 9.8%, and 12.5% for RiboDFM, NA-MPNN, and gRNAde, respectively. This error pattern may be related to RhoDesign’s structural representation, which simultaneously incorporates the coordinates of C1′, N1, and C2 for all four nucleotide types. Although N1 and C2 are present in both purines and pyrimidines, they occupy topologically non-equivalent positions in the two ring systems. Their spatial relationships with C1′ therefore provide readily accessible geometric cues for distinguishing purines from pyrimidines, as reflected by the clearly separated N1-C1′ and C2-C1′ distance distributions shown in Supplementary Fig. 2. Together, these observations suggest that RhoDesign’s high sequence recovery may partly arise from its reliance on simple cues that distinguish purines from pyrimidines, rather than fully capturing the fine-grained sequence– structure constraints needed to preserve native base-pairing patterns and global 3D topology.

Notably, structural self-consistency is positively associated with sequence recovery, but the relationship is far from deterministic (Supplementary Fig. 3). For RiboDFM, sequence recovery showed Spearman correlations of 0.476 with sclDDT and -0.387 with scRMSD, indicating that higher structural self-consistency does not necessarily depend on greater native-sequence recovery. We next compared the agreement with the target structure achieved by predictions from RiboDFM-designed sequences and the corresponding native sequences. As shown in Fig. 2c, many RiboDFM designs achieve sclDDT and scRMSD comparable to or even better than those obtained from native-sequence predictions, despite only partially recovering the native sequence. Similar cases are observed for the other methods, although less frequently (Supplementary Fig. 4). This suggests that RiboDFM can generate sequences with a stronger predicted propensity to adopt the target backbones than the corresponding native sequences.

Fig. 2d-f presents a representative example from New38 (PDB ID: 12CI; released on 13 May 2026). For this target, the RiboDFM design achieves a sequence recovery of 0.585, lower than that of NA-MPNN (0.695), but exhibits better structural self-consistency, with a scRMSD of 6.0 Å, a sclDDT of 68.8 and a scINF of 0.673, compared with 8.3 Å, 51.6 and 0.511 for NA-MPNN, respectively (Fig. 2d). Notably, within the same prediction pipeline, the structure predicted from the RiboDFM-designed sequence is even closer to the experimental structure than that predicted from the native sequence itself. Inspection of the local structures reveals several compensatory substitutions in the RiboDFM design. In the P2 and P3 stems, five native base pairs are replaced by alternative nucleotide pairs while preserving the corresponding pairing relationships in the predicted structure (Fig. 2e). Among the key interactions stabilizing the three-way junction (3WJ), all four native contacts examined are reproduced in the RiboDFM prediction, whereas none are recovered in either the NA-MPNN or native-sequence prediction (Fig. 2f). Notably, although the NA-MPNN design retains the native nucleotide identities at C13-G45, C14-G44, and U41-A43, the corresponding interactions are not reproduced. Together, these observations suggest that RiboDFM captures coupled sequence–structure constraints that permit non-native yet structurally compatible substitutions, thereby facilitating reconstruction of the target fold.

During preparation of this manuscript, AlignIF[12], a recently developed RNA design method incorporating multiple structure alignments (MStA), was reported. We further compared RiboDFM with AlignIF. To distinguish the contribution of structural retrieval, we evaluated both the default AlignIF pipeline and a variant without MStA retrieval, denoted AlignIF (no MStA). The latter uses only the target backbone, providing a matched-input comparison with RiboDFM. The comparison was performed on the aforementioned CASP16 and New38 benchmarks, neither of which temporally overlaps with the AlignIF training data, using the same evaluation protocol with ten different random seeds for each target.

As shown in Supplementary Table 2, RiboDFM matches or exceeds both AlignIF variants in sequence recovery across the evaluated settings. Under the Average setting, RiboDFM achieves 3D structural self-consistency broadly comparable to that of MStA-augmented AlignIF, while its advantage over AlignIF (no MStA) is particularly evident on the New38 dataset. More notably, the advantage of RiboDFM is most pronounced under the Top1 evaluation, which reflects the quality of the most structurally consistent candidate generated within a fixed sampling budget (Supplementary Fig. 5 and Supplementary Table 2). Across both datasets, the Top1 designs by RiboDFM have higher scTM than those generated by either AlignIF variant. On New38, RiboDFM further outperforms both variants across all self-consistency metrics. Specifically, RiboDFM improves Top1 scTM by approximately 0.7–2.2% over AlignIF and 4.3–10.4% over AlignIF (no MStA). These results indicate that, while RiboDFM and MStA-augmented AlignIF achieve comparable average design quality, RiboDFM is more effective at generating high-quality candidates whose predicted structures closely match the target backbones. This stronger Top1 performance may reflect the enhanced sequence-space exploration enabled by RiboDFM’s discrete flow-matching formulation.

### RiboDFM improves structural fidelity in RNA aptamer design

RNA aptamers play important roles in therapeutics, biomolecular detection and fluorescence imaging, making them a key target for RNA design[7, 8, 12]. To evaluate RiboDFM on RNA aptamers, we constructed Apt35, a benchmark comprising 35 experimentally determined RNA aptamer structures that were filtered to remove sequence redundancy with the RiboDFM training set (Methods). Consistent with the previous benchmark, each of the four methods (RiboDFM, NA-MPNN, gRNAde, and RhoDesign) generated ten sequences for every target backbone using the same sampling protocol, and the resulting designs were evaluated at the sequence, and 3D structure levels.

Notably, although all targets in Apt35 were filtered to remove redundancy with the RiboDFM training set, 29 and 24 of the 35 targets overlap with the training sets of NA-MPNN and gRNAde, respectively. Despite this potential advantage for the baseline methods, RiboDFM still achieves strong 3D structure self-consistency on Apt35 (Fig. 3a and Supplementary Table 1). RiboDFM also achieves slightly higher sclDDT and scINF than NA-MPNN (71.4 vs. 71.3 and 0.683 vs. 0.680, respectively), indicating better preservation of local structural geometry and interaction networks. In contrast, NA-MPNN performs slightly better in sequence recovery and the more global 3D consistency metric scTM. These results suggest that RiboDFM’s main advantage on Apt35 lies in recovering structurally important base-pairing and local tertiary interactions, rather than simply maximizing native-sequence recovery or global structural similarity.

**Figure 3.**
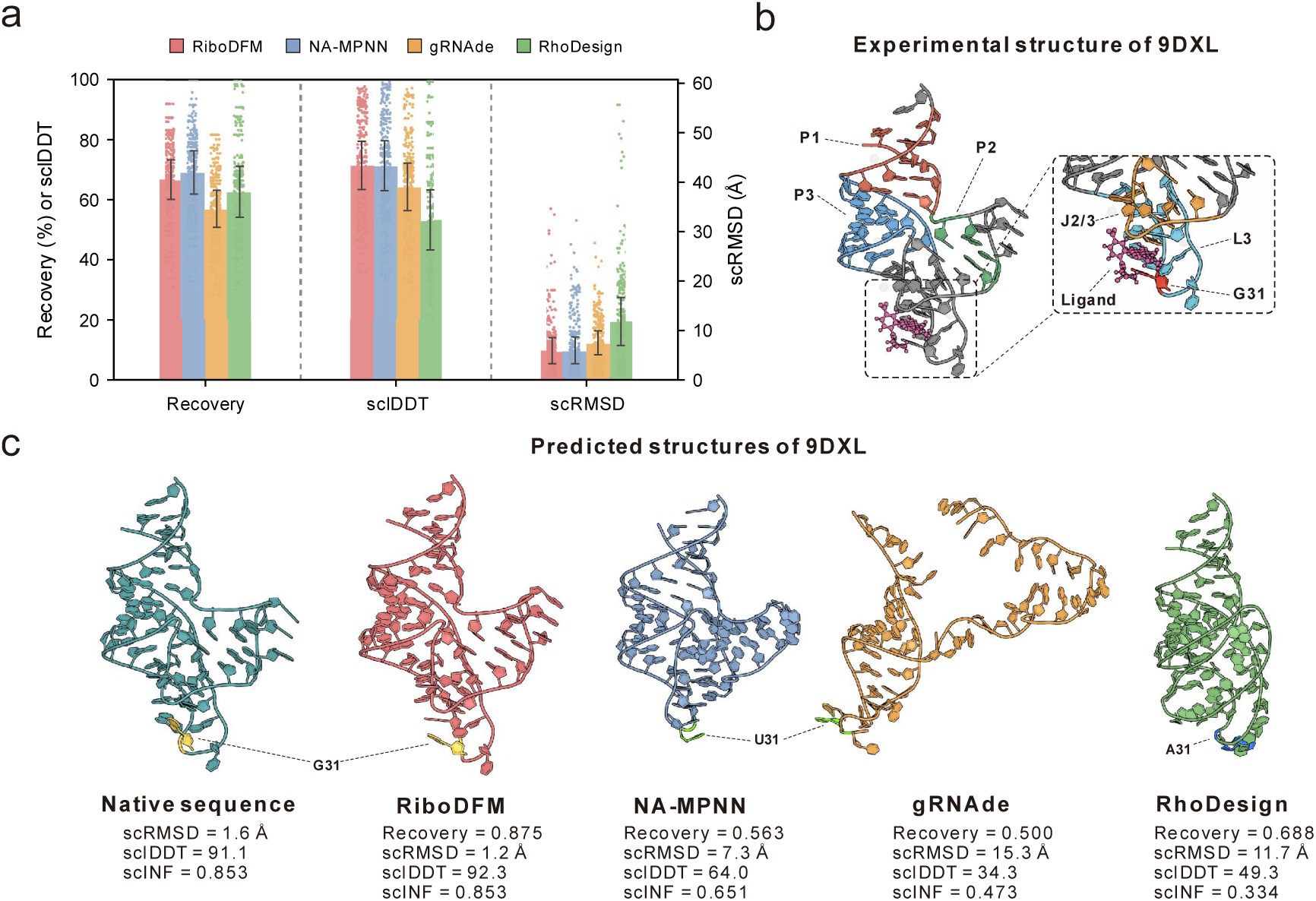
Performance of RiboDFM in RNA aptamer design. **a.** Sequence recovery, sclDDT, and scRMSD on the Apt35 benchmark. Points represent individual designs from ten samples per target, bars indicate mean values, and error bars denote 50% of the standard deviation. Note that a substantial proportion of the targets in this benchmark overlap with the training sets of NA-MPNN (29/35) and gRNAde (24/35). **b, c,** Representative RhoBAST case (PDB ID: 9DXL). **b,** Experimental structure of RhoBAST in complex with MaP555, highlighting the four-way junction architecture and ligand-binding pocket. **c,** Structures predicted from the native sequence and from the scRMSD-top1 design of each method. MaP555 and the functionally critical position 31 (G31 in the native sequence) are highlighted.

We next examined whether RiboDFM can recover the localized structural constraints crucial for aptamer function, using an aptamer target with a chemically well-characterized ligand-binding pocket as a representative case. We illustrate this using the RNA RhoBAST (PDB ID: 9DXL, Fig. 3b) [33]. This is a fluorogenic RNA aptamer crystallized in complex with MaP555, an analogue of the spirocyclic rhodamine dye SpyRho555. It was shown that G31 in RhoBAST plays a critical role and the G31A and G31C substitutions reduce the apparent fluorescence-based affinity for SpyRho555 by more than 430-fold and 640-fold, respectively.

Among the ten designed sequences by RiboDFM, the Top1 design recovers 87.5% of the native sequence and the predicted structure model has scRMSD of 1.2 Å (Fig. 3c), comparable with the model predicted from the native sequence (1.6 Å). This high structural consistency is also observed across the full set of RiboDFM designs: all ten sequences achieve recoveries of 83.3-87.5%, while their predicted structures exhibit scRMSDs of only 1.2-1.5 Å (Supplementary Table 3). The structures predicted from RiboDFM-designed sequences reproduce both the global four-way junction and the relative arrangement of J2/3 and L3 that defines the ligand-binding cleft. In contrast, the lowest scRMSDs achieved by NA-MPNN, gRNAde, and RhoDesign designs are 7.3, 15.3, and 11.7 Å, respectively.

More importantly, sequence-conservation analysis shows that all ten RiboDFM designs retain the functionally critical G31 residue, while U33, A16, and A34 are also strongly conserved (Supplementary Fig. 6). In contrast, G31 is not consistently preserved by the other methods. At position 29, RiboDFM consistently favors G over the native A, representing a purine-preserving substitution within the ligand-binding platform. Rosetta-based energetic evaluation further suggests that the A29G substitution maintains a ligand-interface score comparable to that of the native A29 configuration (−19.555 vs. -19.805 REU; Supplementary Text 1). This case illustrates the potential of RiboDFM to identify non-native sequence solutions that preserve key structural and functional determinants of RNA aptamers.

### RiboDFM enables zero-shot quality assessment of predicted RNA structural models

Beyond sequence generation, the sequence-structure relationship learned by RiboDFM provides a natural signal for reference-free RNA structure quality assessment (QA). A more accurate candidate structure is expected to lead RiboDFM to assign a higher probability to the corresponding native sequence. Based on this principle, we repurposed the pretrained RiboDFM model, without additional training or fine-tuning, to score and rank predicted RNA structural models.

For each candidate model, RiboDFM was conditioned on its backbone and evaluated against the native sequence using a teacher-forced flow trajectory. The probabilities assigned to the correct target nucleotides along the trajectory were aggregated into a global sequence-structure compatibility score, with higher scores indicating better compatibility between the candidate backbone and the native sequence (Methods). We evaluated this zero-shot QA strategy on 6,678 candidate models from 42 CASP16 targets and compared it with eight established RNA QA methods, including three supervised approaches: RNArank[34], lociPARSE[35] and RNA3DCNN[36], and five statistical potentials: DFIRE[37], cgRNASP, cgRNASP-C, cgRNASP-PC[38] and rsRNASP[39]. Ranking performance was assessed using the Spearman correlation between predicted QA scores and lDDT within each target, together with the lDDT of the top-ranked candidate (i.e., Top-1 lDDT).

We first examined whether RiboDFM yields meaningful QA scores for ranking candidate structures. As shown in Fig. 4a, RiboDFM achieves a mean target-wise Spearman correlation of 0.609 between predicted QA scores and ground-truth lDDT, outperforming all compared methods, including lociPARSE (0.597), rsRNASP (0.553), RNArank (0.549), DFIRE (0.540) and the remaining methods (<0.52). At the individual-target level, RiboDFM also achieves higher score-lDDT correlations than RNArank and lociPARSE on 29 and 26 of the 42 targets, respectively (Supplementary Fig. 7). These results demonstrate that RiboDFM can reliably distinguish candidate structures of varying quality without any QA-specific training or fine-tuning.

**Figure 4.**
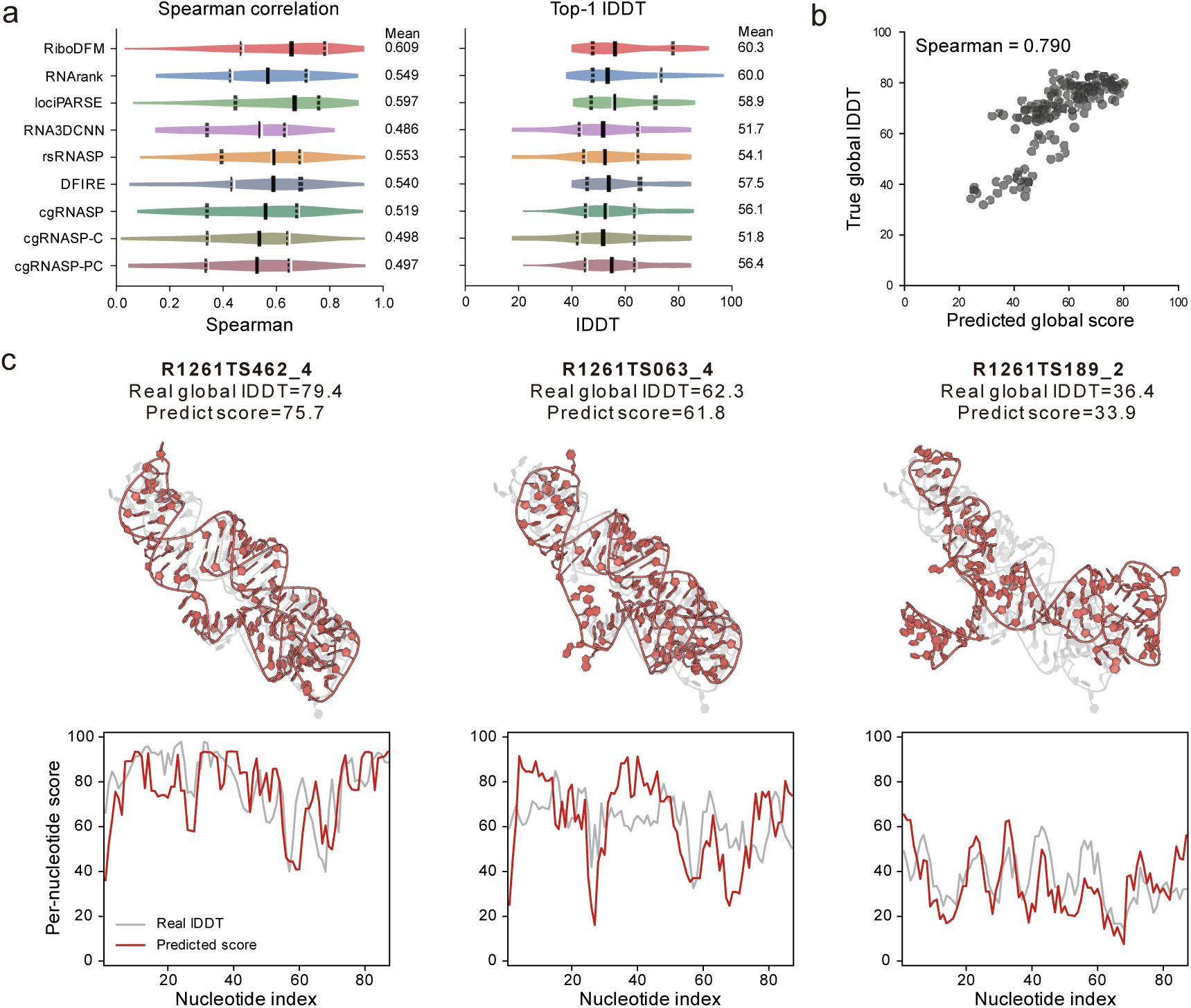
Zero-shot RNA structure quality assessment by RiboDFM. **a**, Distributions of target-wise Spearman correlations and Top-1 lDDT across 42 CASP16 targets for RiboDFM and competing QA methods. Mean values are indicated alongside each distribution. **b,** Correlation between RiboDFM-predicted global QA scores and ground-truth global lDDT values for candidate models of the representative target R1261. **c,** Structural overlays and residue-level quality profiles for three representative R1261 models spanning high, intermediate, and low global quality. In the structural overlays, the native structure is shown in gray and the candidate model in red. In the residue-level profiles, ground-truth local lDDT is shown in gray and the RiboDFM-predicted local QA score in red.

We further evaluated Top-1 model selection performance. For each target, the candidate receiving the highest score from a given QA method was selected, and its ground-truth lDDT was taken as the Top-1 score for that target. The reported Top-1 lDDT was then averaged across the 42 targets. As shown in Fig. 4a, despite being applied in a zero-shot manner, RiboDFM achieves an average Top-1 lDDT of 60.3, closely matching RNArank (60.0) and outperforming lociPARSE (58.9), DFIRE (57.5), and the remaining QA methods (<57). These results indicate that the zero-shot RiboDFM score can be effectively applied to model selection in RNA structure prediction.

The quality-assessment capability of RiboDFM is further illustrated by the representative CASP16 target R1261 (Fig. 4b, c). Across its candidate models, RiboDFM scores show a strong correlation with ground-truth global lDDT (Spearman’s *ρ*=0.790; Fig. 4b), indicating that RiboDFM can distinguish models spanning a broad range of structural quality. We further examined three representative models with high (79.4), medium (62.3), and low (36.4) global lDDT values (Fig. 4c). Their RiboDFM scores are highly consistent with the actual structure qualities, while the residue-level score profiles also broadly recapitulate local variations in lDDT along the RNA chain. These observations suggest that RiboDFM captures not only the relative global quality of candidate structures but also local differences in model accuracy.

In summary, these findings establish the RiboDFM-derived compatibility score as a powerful zero-shot signal for target-specific RNA structure quality assessment. More broadly, the successful transfer from structure-based RNA sequence design to structure evaluation suggests that RiboDFM learns a generalizable representation of RNA sequence–structure compatibility rather than merely a task-specific sequence-generation rule.

### RiboDFM benefits from flow-matching generation, complementary structural training data and sequence-structure priors

To investigate the contributions of sequence-generation strategy, different training-data sources and sequence-structure priors, we retrained four RiboDFM variants from scratch under otherwise identical settings. Specifically, the training data comprise experimentally determined RNA structures from the PDB and large-scale predicted structures for RNAs lacking experimental models (Methods), while the priors include pretrained RNA-FM representations and predicted secondary-structure information. Correspondingly, the four variants were: (1) Autoregressive: replacing the flow-matching sequence-generation procedure with autoregressive decoding while retaining the same training data; (2) Experimental-only: training exclusively on experimentally determined structures; (3) w/o SS head: removing the secondary-structure head; and (4) w/o RNA-FM: replacing the pretrained RNA-FM embeddings with randomly initialized nucleotide embeddings learned from scratch. All variants were evaluated on the same four test datasets using the same ten random seeds as the full RiboDFM model.

We first evaluated the effect of the sequence-generation strategy (Fig. 5a and Supplementary Table 4). Among the four ablation variants, the Autoregressive variant shows the largest overall decline in design performance relative to the full RiboDFM model. Under the Average setting, sequence recovery decreases by 0.017-0.072 across the four test sets. Structural self-consistency also deteriorates overall, with sclDDT decreasing by 0.6-4.5, consistent reductions in scTM and scINF, and increased scRMSD on three of the four datasets. A similar overall trend is observed under the Top1 setting, particularly on New38, where sclDDT, scTM and scINF decrease by 3.1, 0.031 and 0.038, respectively, and scRMSD increases by 0.6 Å. These results support the advantage of discrete flow matching over autoregressive decoding in improving sequence recovery and overall structural self-consistency.

**Figure 5.**
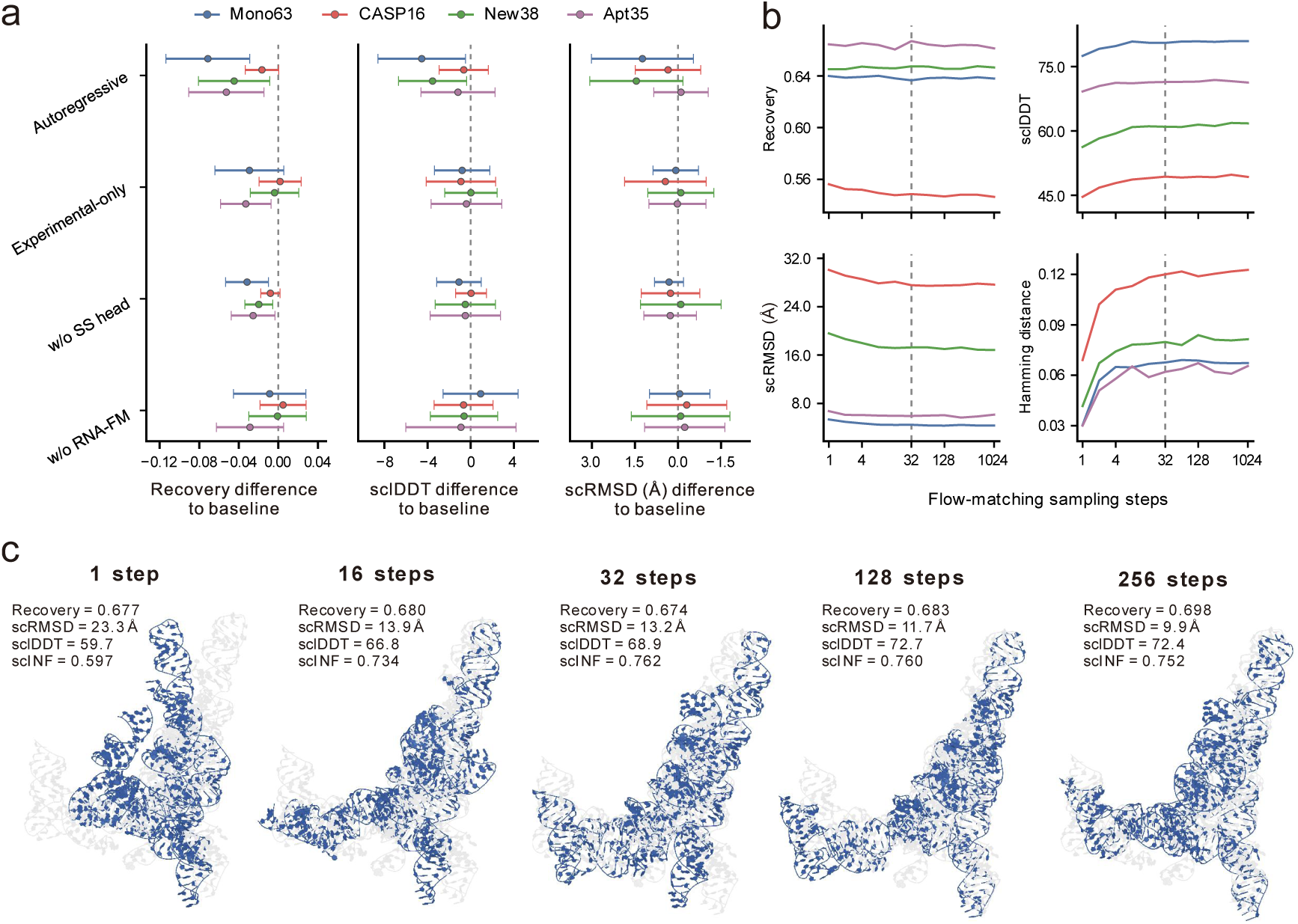
Ablation and sampling-step analyses of RiboDFM. **a**, Performance changes of four RiboDFM variants relative to the full model in sequence recovery, sclDDT, and scRMSD across the Mono63, CASP16, New38, and Apt35 benchmarks. Points indicate the mean paired target-level differences, and error bars represent 50% of the standard deviation across paired targets. **b,** Effects of the number of flow-matching sampling steps on sequence recovery, sclDDT, scRMSD, and sequence diversity measured by mean pairwise Hamming distance. The dashed line marks the default 32-step setting. **c,** Representative example for target 9K9G showing performance changes as the number of sampling steps increases. For each setting, the scRMSD-top1 design among ten generated sequences is shown.

We next examined the contribution of predicted training structures (Fig. 5a and Supplementary Table 4). The Experimental-only variant remains close to the full RiboDFM model on CASP16 and New38, but shows decreases of 0.009–0.033 in average sequence recovery and scINF on Mono63 and Apt35. Under the Top1 setting, sequence recovery shows slightly larger decreases of 0.032 and 0.035 on Mono63 and Apt35, respectively, while scINF remains lower across all four datasets. This indicates that augmenting the limited experimentally determined RNA structures in the PDB with large-scale predicted structures provides additional benefit, potentially by improving coverage of structural and sequence diversity.

Finally, we evaluated the contributions of the sequence–structure priors provided by the secondary-structure head and RNA-FM (Fig. 5a and Supplementary Table 4). Under the Average setting, removing the secondary-structure head consistently reduces sequence recovery across all four test sets, with decreases ranging from 0.008 to 0.032. Its effects on structural self-consistency are more modest, with changes of up to 1.1 in sclDDT and 0.3 Å in scRMSD. The contribution of RNA-FM is more dataset-dependent. Its removal has the strongest effect on Apt35, decreasing sequence recovery from 0.667 to 0.638, sclDDT from 71.4 to 70.5, and scINF from 0.683 to 0.659, while producing smaller changes on the other three test sets. Under the Top1 setting, both components have a more pronounced contribution on Apt35, where removing the secondary-structure head or RNA-FM decreases sequence recovery from 0.676 to 0.645 or 0.649, respectively. Overall, the full RiboDFM model, which integrates backbone-derived secondary-structure information with pretrained sequence representations, provides the strongest and most consistent performance among the evaluated variants across the four benchmark datasets.

### RiboDFM balances design quality and computational efficiency through adjustable sampling steps

Beyond the default 32-step decoding scheme, the discrete flow-matching framework provides substantial flexibility for sequence generation. As illustrated in Fig. 1c, unlike autoregressive approaches that sequentially predict nucleotides one position at a time, RiboDFM predicts nucleotide probability distributions for all residues at each flow step and updates the sequence state in parallel (Methods). This formulation allows the number of sampling steps to be adjusted at inference, providing a tunable trade-off between computational efficiency and iterative sequence refinement. We therefore systematically examined how the number of sampling steps affects design performance. Specifically, we ran RiboDFM using flow trajectories from *t* = 0 to *t* = 1 discretized into 1 to 1,024 sampling steps. The corresponding results across four benchmark datasets are summarized in Fig. 5b and Supplementary Table 5.

Interestingly, RiboDFM achieves high sequence recovery even with single-step sampling, and increasing the number of sampling steps has only a limited effect on this metric. In contrast, 3D structural self-consistency generally improves as more sampling steps are used. On the CASP16 dataset, for example, one-step sampling yields the highest average sequence recovery of 0.556, whereas the default 32-step setting produces a slightly lower value of 0.549. However, sclDDT substantially improves from 44.6 to 49.3, accompanied by an increase in scINF from 0.420 to 0.543. This discrepancy is consistent with our observation that sequence recovery and structural self-consistency are not tightly coupled (Supplementary Fig. 3). In addition, sequence diversity generally increases with the number of sampling steps, as reflected by the higher mean pairwise Hamming distance (Fig. 5b). These results suggest that additional flow-matching steps primarily enhance compatibility with the target structure and broaden exploration of sequence space, rather than driving the designed sequences closer to the native sequence.

Meanwhile, we found that most gains in structural self-consistency occur at relatively low numbers of sampling steps, after which performance gradually plateaus. Across the four datasets, increasing the number of steps from the default 32 to 1,024 changes the average sclDDT by at most 0.8 points and the average scRMSD by at most 0.5 Å, while increasing the inference time from 1.6–3.6 to 44.1–58.2 s per sequence (Supplementary Fig. 8). Nevertheless, additional sampling steps can still benefit individual targets, as illustrated by 9K9G (Fig. 5c). Although sequence recovery remains broadly stable across different sampling steps, structural self-consistency improves substantially with additional steps. The Top1 scRMSD decreases from 23.3 Å with one-step sampling to 13.2 Å with 32 steps and 9.9 Å with 256 steps, while sclDDT increases from 59.7 to 68.9 and 72.4, respectively.

Overall, the default 32-step RiboDFM setting achieves near-optimal performance with a relatively short average inference time of approximately 2.3 s per sequence. Its performance can be further adjusted by varying the number of sampling steps: fewer steps enable faster generation for high-throughput applications with minimal loss in sequence recovery, whereas additional steps may further improve structural self-consistency for specific RNA targets. This flexibility arises from the parallel sequence refinement enabled by discrete flow matching, in contrast to the position-by-position decoding used by conventional autoregressive models.

### Probability consistency along the flow trajectory provides a model-internal confidence measure for designed sequences

To estimate design quality directly from the model without relying on computationally expensive structural self-consistency evaluation, we leveraged the consistency of nucleotide probabilities during the late stages of the flow trajectory. A well-supported design should receive consistently high probabilities for its nucleotide choices across successive sampling steps, rather than only at the terminal state. We therefore defined a sequence-level confidence score based on the final *K_c_* flow steps, with *K_c_*=10 used in this study. At each of these steps, we recorded, for every residue, the probability assigned to the nucleotide present in the final generated sequence and calculated the geometric mean over all residues and steps:

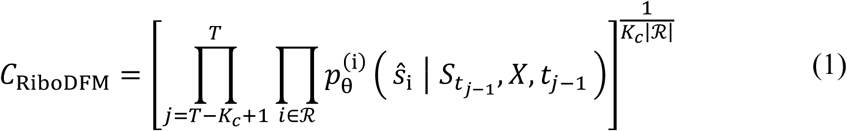

where *^S_i_* represents the nucleotide at position *i* in the final generated sequence; *S_t_* is the intermediate sequence state at flow step *t*_1_; *X* denotes the target backbone; *R* the set of residues; and *T* is the total number of sampling steps. A higher *C* score indicates that RiboDFM consistently assigns high probabilities to the final nucleotide choices during the late stages of generation. We therefore used *C* as a model-internal confidence estimate for each designed sequence and examined whether it reflects structural self-consistency.

We examined the relationships of *C*_RiboDFM_ with sequence recovery, sclDDT and scRMSD on Mono63, CASP16 and New38. Although *C*_RiboDFM_ is less strongly correlated with sequence recovery than the confidence score reported by NA-MPNN (denoted *C*_NA-MPNN_), it shows stronger correlations with structural self-consistency, especially on the more challenging CASP16 and New38 datasets (Supplementary Fig. 9). On the CASP16 dataset, the Spearman correlations of *C*_RiboDFM_ with sclDDT and scRMSD are 0.594 and -0.749, respectively, substantially stronger than those of *C*_NA-MPNN_ (0.212 and -0.416). Similarly, on New38, *C*_RiboDFM_ also correlates more strongly with sclDDT (0.633 vs. 0.496) and scRMSD (−0.608 vs. -0.482). Notably, on both CASP16 and New38, *C*_RiboDFM_ correlates more strongly with the structural metrics than with sequence recovery, suggesting that it captures sequence–structure compatibility beyond native-sequence recovery and can serve as a model-internal proxy for structural self-consistency. This advantage highlights a distinctive benefit of the flow-matching framework, in which probability consistency across successive states along the generation trajectory provides a richer confidence signal than a single-step autoregressive prediction.

## Discussion

In this study, we developed RiboDFM, a backbone-conditioned discrete flow-matching framework for RNA sequence design. Across complementary benchmarks, RiboDFM achieved strong native-sequence recovery and favorable tertiary-structure self-consistency, including on recently released structures and aptamer-focused targets. RiboDFM also highlights several advantages of discrete flow matching as a sequence-generation framework. Unlike conventional autoregressive approaches that perform sequential, one-by-one generations, RiboDFM predicts nucleotide distributions for all positions and iteratively refines multiple positions in parallel. A single sampling step already yields high sequence recovery, whereas additional steps primarily improve structural self-consistency and sequence diversity. This behavior suggests that iterative flow refinement helps coordinate nucleotide choices across positions to achieve globally more structure-compatible solutions, rather than simply driving the generated sequence toward the native sequence.

Interestingly, the flow trajectory also provides informative signals beyond the final generated sequence. Designs that remain consistently supported across successive late-stage flow states tend to exhibit better structural self-consistency. Accordingly, the trajectory-derived confidence score correlates more strongly with structural self-consistency and more reliably prioritizes high-quality designs than the autoregressive confidence score evaluated in this study. The learned sequence-structure compatibility signal further enables RiboDFM to rank predicted RNA structural models without quality-assessment-specific training. These findings indicate that the intermediate flow states are not merely computational waypoints in sequence generation. Instead, the probability patterns along the trajectory provide interpretable signals for confidence estimation and downstream structural model assessment. Overall, our findings establish RiboDFM as an effective and flexible framework for 3D RNA structure-based sequence design and demonstrate that sequence-design models can learn transferable information about RNA sequence-structure compatibility.

Despite its strong performance, RiboDFM has several limitations. First, our structural self-consistency metrics rely on computational predictions. Given the inherent inaccuracies of current RNA structure prediction, experimental validation remains the gold standard; while this falls outside the scope of our study, we have open-sourced all code and data to enable community-driven application and wet-lab validation. Second, RiboDFM does not explicitly model interacting partners, such as proteins and ligands, and therefore cannot directly optimize intermolecular recognition. Third, RiboDFM conditions on a single static backbone, whereas many RNAs populate heterogeneous conformational ensembles or undergo functionally important structural transitions. Structural homologs may also provide useful complementary information, as demonstrated by the multiple-structure-alignment strategy introduced in AlignIF[12]. Future work integrating explicit binding partners, conformational ensembles, evolutionary homologs, and experimental constraints will be crucial for the robust design of functionally specialized RNAs.

## Methods

### RiboDFM algorithm

RiboDFM is a backbone-conditioned discrete flow-matching framework consisting of three learnable modules: a geometric backbone encoder, an auxiliary secondary-structure prediction head, and a flow-matching decoder. The first two modules derive conditioning representations from the target backbone, while the flow-matching decoder predicts position-wise nucleotide distributions conditioned on these backbone representations and the current sequence state. During inference, a discrete flow sampler uses the decoder-predicted nucleotide distributions to iteratively update the sequence state along a discretized flow trajectory, thereby progressively transforming a fully masked sequence into a complete RNA sequence compatible with the target backbone. The individual components and procedures are described in detail below.

#### Backbone encoder

Each RNA residue is represented as a node in a residue-level *k*-nearest-neighbor (*k*-NN) graph, with edges connecting its *k* nearest neighbors (*k* = min(128, *L* − 1); *L* is the sequence length) according to their C4′-C4′ distances. Each residue is represented by three atomic positions: C4′, C1′, and a virtual base-nitrogen atom reconstructed from the C4′, C1′, and C2′ coordinates (inspired by NA-MPNN[11]). For each directed edge ( *i* → *j* ), the nine pairwise distances between the three representative atoms of residues *i* and *j* are expanded using 16 radial basis functions spanning 0–20 Å. The resulting distance features are concatenated with an embedding of the relative sequence-index offset (*j* − *i* ). These features are projected and normalized to initialize the edge representations, whereas the node representations are initialized to zero. The node and edge representations are subsequently refined through three stacked message-passing layers with independent parameters. Each layer employs gated multilayer perceptrons, residual connections, and layer normalization.

#### Secondary-structure prediction head

To provide an explicit base-pairing prior derived solely from the target backbone, the encoded edge representations are passed through a two-layer multilayer perceptron to predict a binary base-pairing probability for each *k*-nearest-neighbor edge. The predicted probabilities are incorporated into the flow-matching decoder through a learnable gating function that modulates message passing along structural edges, thereby allowing edges with higher predicted base-pair probabilities to contribute more strongly. During training, the decoder is conditioned, with equal probability, on either the predicted pairing probabilities or the available ground-truth base-pair labels, whereas only the predicted probabilities are used during inference. The secondary-structure head is optimized using an auxiliary binary focal loss. A stop-gradient operation is applied before the predicted pairing probabilities are passed to the decoder, such that gradients from the flow-matching decoder are not propagated through this head.

#### Flow-matching decoder

At each flow step *t*, the current sequence state *S_t_* is denoted by

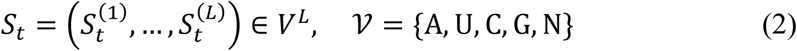

where N denotes an unresolved nucleotide, and *L* is the sequence length. The current sequence state is encoded using the 12-layer RNA-FM model, and the 640-dimensional representations from its final layer are projected to the hidden dimension of RiboDFM (256-dimensional). A Fourier embedding of the flow time *t* is added to both the projected RNA-FM representations and the backbone node representations. The resulting sequence representations, backbone node and edge representations, and predicted base-pairing probabilities are then jointly processed by the flow-matching decoder, which consists of three message-passing layers with independent parameters.

For each residue *i* , the decoder predicts a categorical distribution over the four canonical nucleotide types,

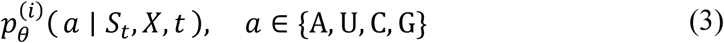

Although the mask token N may occur in the input sequence state, the decoder predicts probabilities only over the four canonical nucleotides. These predicted distributions provide the nucleotide preferences used by the discrete flow sampler to update the sequence state (introduced below).

#### Discrete probability path and training

Given a native sequence *S*_1_ , RiboDFM defines a discrete probability path connecting the fully masked state *S* = (N, … , N) to the native sequence *S*_1_. At each training iteration, a continuous flow time *t*∼*Ʋ*(0,1) is sampled. One of four interpolation schedules is then selected uniformly:

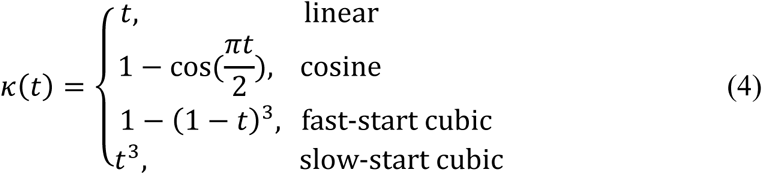

The intermediate sequence state *S_t_* is constructed using a binary mask *M_t_*, whose entries are sampled independently as *M_t_*^(*i*)^ ∼ Bernoulli (*k*(*t*)):

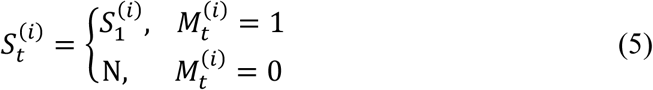

Thus, each native nucleotide is retained with probability *κ*(*t*) , whereas the remaining positions are replaced by the mask token N . The expected fraction of resolved positions therefore increases continuously from 0 at *t*=0 to 1 at *t*=1. Conditioned on the target backbone *X*, the partially masked sequence *S_t_*, and the flow time *t*, RiboDFM is trained to directly predict the complete native sequence *S*_1_ in a single flow step. The complete training procedure is summarized in Supplementary Algorithm 1.

#### Discrete flow sampler

The discrete flow sampler converts the decoder-predicted nucleotide distributions into iterative sequence-state updates along a discretized flow trajectory. During inference, the sequence is initialized to the fully masked state *S* = (N, … , N). The continuous flow trajectory from *t* = 0 to *t* = 1 is discretized into a predefined number of steps (32 steps by default). At each step *k*, the decoder predicts a nucleotide distribution for every position in parallel.

To distinguish a categorical sequence state from a probability distribution, let

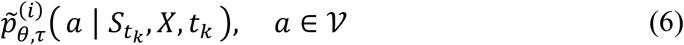

denote the temperature-scaled decoder distribution. The probabilities over the four canonical nucleotides were obtained by applying temperature scaling to the decoder logits before softmax, that is, Softmax 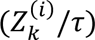, where 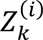 denotes the decoder logits for residue *i* at step *k*. Zero probability was assigned to the mask token: 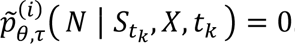. The transition distribution for position *i* is defined as

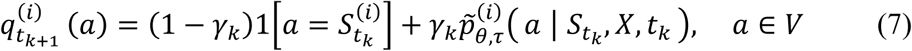

followed by categorical sampling:

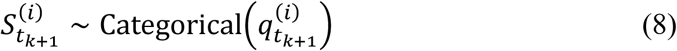

Eq. (7) defines a mixture between retaining the current nucleotide state and resampling from the decoder-predicted distribution, with their relative contributions controlled by the transition weight 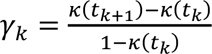. Equivalently, each position can be viewed as independently retaining its current state with probability 1 − γ*_k_* or being resampled from the decoder distribution with probability γ*_k_*. Repeating this procedure progressively transforms the fully masked initial state into the final designed RNA sequence. Unless otherwise specified, the cosine schedule was used during inference. The complete sampling and confidence-estimation procedures are summarized in Supplementary Algorithm 2.

### Construction of datasets

#### Test sets

Four benchmark datasets were used to evaluate RNA sequence design performance: Mono63, CASP16, Apt35 and New38. Mono63 was adopted from the NA-MPNN test set and contains 63 monomeric RNA structures. The initial CASP16 set contained 31 RNA targets from the CASP16 experiment. Six targets sharing ≥80% sequence identity with the NA-MPNN training or validation sets were removed using CD-HIT-EST[40], resulting in 25 targets. Apt35 consists of 35 nonredundant, experimentally determined RNA aptamer structures collected from the PDB. New38 was constructed to assess temporal generalization and contains 38 monomeric RNA structures released in the PDB between 21 January 2025 and 1 June 2026, after the training-data cutoffs of the evaluated design methods.

#### Training set from PDB

We collected all RNA-containing structures released in the PDB before 21 January 2025 and extracted their RNA components. Multichain structures were first separated and subsequently regrouped into individual samples according to interchain base-pairing interactions, following the data-processing procedure used in trRosettaRNA2. This procedure yielded 12,948 RNA samples (representing an expanded version of the trRosettaRNA2 training set). To avoid data leakage, sequences with similarity to Mono63, CASP16, or Apt35 were removed using CD-HIT-EST with the parameters -c 0.80 -n 5 -G 1 -s2 0.5 -S2 100000 -aS 0.80 -aL 0.5 -mask N -g 1 -p 1 -d 0 -T 16 -M 0. The remaining sequences were clustered at 80% sequence identity, producing 2,469 clusters. Of these, 2,419 clusters containing 12,036 RNA samples were used for training, and 50 clusters containing 140 samples were reserved for validation.

#### trRosettaRNA2-predicted training data

The PDB-derived training set was supplemented with structures predicted by trRosettaRNA2 from bpRNA and RNAcentral sequences. For bpRNA sequences, the annotated secondary structures were provided as input for structure prediction. For RNAcentral, only predictions with pLDDT >60 were retained. Sequences sharing ≥80% identity with any test target were removed using the same CD-HIT-EST procedure mentioned above. The final predicted training set contained 13,479 bpRNA-derived and 136,349 RNAcentral-derived structures.

### Loss function and training procedure

Given an intermediate sequence *S_t_* sampled according to Eq. (5), RiboDFM was trained to recover the corresponding native sequence *S_1_*. The total training loss was defined as

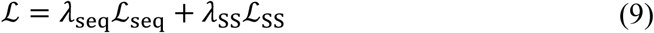

where ℒ_seq_ is the cross-entropy loss between the predicted nucleotide distributions and native nucleotide identities, and ℒ is the binary focal loss for base-pair prediction on graph edges. The focal-loss parameters were set to *α*=0.8 and *γ*=2.0, with *λ*_seq_=1.0 and *λ*_SS_=10.0. Label smoothing of 0.1 was applied to ℒ, and residues with unknown nucleotide identities were excluded from the loss calculation.

RiboDFM was optimized using AdamW with an initial learning rate of 5e-4 and a weight decay of 0.01. The learning rate was reduced by a factor of 0.5 if the validation loss did not improve for 20 consecutive epochs. The pretrained RNA-FM model was frozen. At each epoch, all PDB-derived structures and a randomly sampled 5% subset of the trRosettaRNA2-predicted structures were used for training. Two models with identical architectures were trained, and their predicted nucleotide probabilities were averaged during sampling. The model checkpoint with the highest sequence recovery on the validation set was selected for evaluation.

### Baseline methods

RiboDFM was compared with four representative 3D backbone-conditioned RNA sequence design methods: RhoDesign, gRNAde, NA-MPNN and AlignIF. These methods differ mainly in how the target RNA structure is represented and encoded. RhoDesign uses nucleotide-level geometric features derived from backbone and base-related atomic coordinates (C4’, C1’, N1, C2, C5’, O5’, P), whereas gRNAde adopts a coarse-grained three-bead representation consisting of P, C4′, and the glycosidic nitrogen atom (N1 for pyrimidines and N9 for purines). AlignIF likewise incorporates the glycosidic nitrogen atom into its geometric features, while further integrating structural information from similar RNA folds through multiple structure alignment (MStA). By comparison, NA-MPNN represents RNA using a more detailed set of sugar–phosphate backbone atoms together with a reconstructed virtual base-N atom.

All baseline methods were run locally using their official implementations and released pretrained models. The same target backbones were provided to all methods. For each target, ten sequences were generated using different random seeds at a sampling temperature of 0.1, with other method-specific settings kept at their defaults. RhoDesign was run without its optional secondary-structure input so that all methods were conditioned only on the target backbone.

AlignIF was evaluated on the CASP16 and New38 datasets, as neither dataset directly overlaps with the training set used by AlignIF, enabling a more appropriate evaluation of its released pretrained model. We considered both the default AlignIF pipeline, which performs MStA retrieval using the structure database provided by the authors, and an ablated setting without MStA retrieval, referred to as AlignIF (no MStA). The latter uses only the target structure, whereas the default model additionally incorporates structurally similar RNAs retrieved by MStA.

### Design performance and structure prediction

Performance was assessed at the sequence and tertiary-structure levels.

### Sequence recovery

Sequence recovery was calculated as the fraction of designed nucleotides identical to the native sequence. Residues with unknown native nucleotide identities were excluded.

### RNA tertiary-structure prediction and consistency

To improve the robustness of structural evaluation and reduce dependence on any single predictor, three-dimensional structures were independently predicted from each designed sequence (without utilizing multiple sequence alignments) using trRosettaRNA2[20], AlphaFold 3[31] and Protenix[32]. **trRosettaRNA2** was run using its default parameters and settings, and the unrefined predicted structure was used for evaluation. **AlphaFold 3** was run independently for each designed sequence using five random seeds and five samples per seed, and the structure with the highest ranking score was retained for evaluation. **Protenix** was run with the Protenix-v1 model using five random seeds and five samples per seed, and the structure achieving the highest reported pLDDT was selected for evaluation. Among the three predictor-specific structures, the one with the lowest C1’-RMSD to the target backbone was selected as the representative structure for each designed sequence, and all subsequent 3D structural self-consistency metrics were calculated using this same structure.

The selected representative predicted structure for each designed sequence was evaluated against the corresponding experimental structure to calculate C1’-based lDDT, TM-score, C1’-RMSD, and interaction network fidelity (INF), denoted sclDDT, scTM, scRMSD, and scINF, respectively. Specifically, C1’-based lDDT was calculated using the RNA-adapted implementation[34], with a 30-Å inclusion radius and distance-error thresholds of 1, 2, 4, and 6 Å. TM-score and RMSD were calculated using US-align[41] in RNA mode with C1’ atoms. INF was calculated by independently identifying base-pair interactions in the predicted and experimental structures using MC-Annotate[42] and taking the geometric mean of base-pair precision and recall[43].

### Confidence scoring

For each RiboDFM-generated sequence, the confidence score was computed based on the tail-trajectory geometric mean formulation described in the Results. During sequence generation, the raw, untempered predicted probability *p*^(*i*)^ assigned to the final sampled nucleotide *^S_i_* was recorded at each of the final *K_c_* = 10 discrete flow steps. Residues with invalid or missing input coordinates were excluded from the set of valid designed residues *R*. For NA-MPNN, the default confidence score reported by its official implementation was used for all baseline comparisons.

### Zero-shot RNA structure quality assessment

RiboDFM was adapted to RNA structure quality assessment without additional training or fine-tuning. In the original sequence-design task, RiboDFM predicts a sequence conditioned on a given backbone. For structure quality assessment, this formulation was reversed: the RNA sequence was fixed, and each candidate structure was evaluated according to how well RiboDFM supported the known sequence when conditioned on that structure.

Given a candidate backbone *X* and its corresponding target sequence *S* = (*s*_1_, … , *s_L_*) , let *R* denote the set of valid residues and let *π* = *π*_1_, … , *π*_|*R*|_ be a randomly sampled permutation of their indices. Starting from a fully masked sequence, target nucleotides were progressively introduced according to this order. At step *m*, the positions (*π*_1_, … , *π_m_*) had already been filled with their target nucleotides, whereas the remaining positions were still masked. For the next residue *i* = *π_m_* , RiboDFM predicted a nucleotide distribution conditioned on the candidate backbone, the current teacher-forced sequence state, and the corresponding flow time. Before *s_i_* was added to the sequence context, its residue-level compatibility score was recorded as

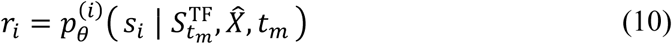

where *p*_0_^(*i*)^ ( ) denotes the nucleotide distribution predicted for residue *i*, and 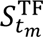 denotes nthe teacher-forced sequence state immediately before residue *i* = *π_m_* was revealed. After *r_i_* was recorded, the target nucleotide *s_i_* was inserted into the sequence state and used as context for subsequent predictions. This one-by-one teacher-forcing trajectory was used specifically for QA and differed from the parallel multistep sampling procedure used for RNA sequence design.

To reduce residue-level noise and incorporate information from the local three-dimensional environment, the compatibility score of each residue was averaged with those of its four nearest valid spatial neighbors. Specifically, *N*(*i*) contains residue *i* and its four nearest valid neighbors according to C4′-C4′ distance. The local QA score was calculated as

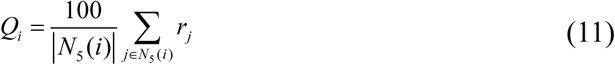

where the factor of 100 expresses the score on a scale from 0 to 100. The global QA score of a candidate structure was then obtained by averaging the local scores over all valid residues:

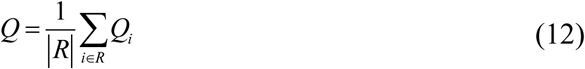

Residues with invalid nucleotide identities or missing structural coordinates were excluded from *R*. A higher *Q* score indicates stronger compatibility between the candidate backbone and its target sequence. Candidate models were ranked separately for each target according to their global *Q* scores. The complete zero-shot structure quality assessment procedure is summarized in Supplementary Algorithm 3.

The QA benchmark comprised 6,678 candidate structural models from 42 CASP16 targets. RiboDFM was compared with RNArank, lociPARSE, RNA3DCNN, DFIRE, cgRNASP, cgRNASP-C, cgRNASP-PC, and rsRNASP. For each target, ranking performance was evaluated using the Spearman rank correlation coefficient between the predicted global QA scores and the corresponding lDDT values. Top-1 performance was defined as the lDDT of the candidate model assigned the highest QA score, and the resulting values were averaged over the 42 targets. Experimental reference structures and lDDT values were used only for benchmark evaluation and were not provided to RiboDFM during scoring. Because RiboDFM was trained exclusively for RNA sequence design without exposure to structural decoys or model-quality labels, the released pretrained model was applied directly to this task.

## Data availability

All training and benchmark datasets are available at: https://yanglab.qd.sdu.edu.cn/RiboDFM/.

## Code availability

The source code is publicly available at GitHub: https://github.com/YangLab-SDU/RiboDFM.

## Author contributions

J.Y. conceived and supervised the project; X.L. designed and implemented the experiments; X.A. participated in model development and experimental design. W.W. and Z.P. co-supervised the study; All authors reviewed and approved the final version of the paper.

## Acknowledgements

This work is supported by the National Natural Science Foundation of China (NSFC T2225007, 32430063, 62501364), and Fundamental Research Funds for the Central Universities.

