## Supplementary Material for "RiboDFM: Robust RNA sequence design using backbone-conditioned discrete flow matching"

### Supplementary Information

#### Supplementary Algorithms

To complement the mathematical description in the Methods, the main computational procedures of RiboDFM are summarized as pseudocode below. Supplementary Algorithms 1–3 describe model training, sequence sampling with confidence estimation, and zero-shot RNA structure quality assessment, respectively.

##### Supplementary Algorithm 1. One training step of RiboDFM.

Supplementary Algorithm 1 summarizes a training step of RiboDFM. A partially masked sequence is constructed at a sampled flow time, and the combined sequence and base-pair prediction loss is used to update the trainable parameters while keeping RNA-FM frozen.

**Supplementary Algorithm 1** One training step of RiboDFM

**Input:** Backbone $X$, native sequence $S_1$, base-pair labels $Y_{\mathrm{SS}}$, valid residues $\mathcal{R}$
**Output:** Updated parameters $\theta$

$\kappa \leftarrow$ RandomSchedule
$t \sim \mathcal{U}(0,1)$
**for** $i \in \mathcal{R}$ **do**
  $M_t^{(i)} \sim \mathrm{Bernoulli}(\kappa(t))$
  $S_t^{(i)} \leftarrow \begin{cases} S_1^{(i)}, & M_t^{(i)} = 1, \\ N, & M_t^{(i)} = 0 \end{cases}$
**end for**
$(H_V, H_E) \leftarrow$ BackboneEncoder$(X)$
$P_{\mathrm{SS}} \leftarrow$ SecondaryStructureHead$(H_E)$
$b \sim \mathrm{Bernoulli}(0.5)$
$Y_{\mathrm{cond}} \leftarrow Y_{\mathrm{SS}}$ if $b = 1$; otherwise stopgrad$(P_{\mathrm{SS}})$
$C_X \leftarrow (H_V, H_E, Y_{\mathrm{cond}})$
$E_t \leftarrow$ RNA-FM$(S_t)$ ▷ RNA-FM is frozen
$Z_t \leftarrow$ FlowMatchingDecoder$(E_t, C_X, t)$
$\mathcal{L}_{\mathrm{seq}} \leftarrow$ CrossEntropy$(Z_t, S_1)$
$\mathcal{L}_{\mathrm{SS}} \leftarrow$ FocalLoss$(P_{\mathrm{SS}}, Y_{\mathrm{SS}})$
$\mathcal{L} \leftarrow \lambda_{\mathrm{seq}}\mathcal{L}_{\mathrm{seq}} + \lambda_{\mathrm{SS}}\mathcal{L}_{\mathrm{SS}}$
$\theta \leftarrow$ AdamWUpdate$(\theta, \nabla_\theta \mathcal{L})$
**return** $\theta$

**Supplementary Algorithm 2. RiboDFM discrete flow sampling and confidence estimation.**

Supplementary Algorithm 2 summarizes the discrete flow sampler used for parallel sequence generation and trajectory-based confidence estimation. Starting from a fully masked sequence, transition distributions for all valid residues are constructed and sampled in parallel at each step. The nucleotide probabilities before temperature scaling from the final $K_c$ steps ($K_c = 10$ by default) are used to calculate the confidence score. Note that confidence estimation is performed only under the default 32-step sampling setting.

---

**Supplementary Algorithm 2** RiboDFM discrete flow sampling and confidence estimation

---

**Input:** Backbone $X$, sequence length $L$, valid residues $\mathcal{R}$, sampling steps $T$, schedule $\kappa$, temperature $\tau$, confidence window $K_c$

**Output:** Designed sequence $\widehat{S}$ and confidence score $C_{\text{RiboDFM}}$

$(H_V, H_E) \leftarrow \textsc{BackboneEncoder}(X)$
$P_{\text{SS}} \leftarrow \textsc{SecondaryStructureHead}(H_E)$
$C_X \leftarrow (H_V, H_E, P_{\text{SS}})$
$S_{t_0} \leftarrow (N, \ldots, N)$
**for** $k = 0$ to $T - 1$ **do**
  $t_k \leftarrow k/T$
  $t_{k+1} \leftarrow (k+1)/T$
  $E_k \leftarrow \text{RNA-FM}(S_{t_k})$
  $Z_k \leftarrow \textsc{FlowMatchingDecoder}(E_k, C_X, t_k)$
  $P_{k+1} \leftarrow \text{Softmax}(Z_k)$ ▷ untempered probabilities
  $\widetilde{P}_{k+1} \leftarrow \textsc{ExtendToVocabulary}(\text{Softmax}(Z_k/\tau),\ P(N) = 0)$
  $\gamma_k \leftarrow \dfrac{\kappa(t_{k+1}) - \kappa(t_k)}{1 - \kappa(t_k)}$
  **for** $i \in \mathcal{R}$ **in parallel do**
    $q^{(i)}_{t_{k+1}}(a) \leftarrow (1 - \gamma_k)\mathbf{1}[a = S^{(i)}_{t_k}] + \gamma_k \widetilde{P}^{(i)}_{k+1}(a)$
    $S^{(i)}_{t_{k+1}} \sim \text{Categorical}\left(q^{(i)}_{t_{k+1}}\right)$
  **end for**
**end for**
$\widehat{S} \leftarrow S_{t_T}$
**if** $N \in \widehat{S}$ **then**
  $\widehat{S} \leftarrow \textsc{FillResidualMasks}(\widehat{S}, \widetilde{P}_T)$
**end if**
$C_{\text{RiboDFM}} \leftarrow \left[\prod_{j=T-K_c+1}^{T} \prod_{i \in \mathcal{R}} P^{(i)}_j(\widehat{s}_i)\right]^{1/(K_c|\mathcal{R}|)}$
**return** $\widehat{S}$, $C_{\text{RiboDFM}}$

---

**Supplementary Algorithm 3. Zero-shot RNA structure quality assessment by RiboDFM.**

Supplementary Algorithm 3 summarizes zero-shot RNA structure quality assessment. Native nucleotides are revealed in a random order, and the probabilities assigned immediately before each reveal are aggregated over local C4′-based neighborhoods and all valid residues to obtain residue-level and global quality scores.

---

**Supplementary Algorithm 3** Zero-shot RNA structure quality assessment

---

**Input:** Predicted backbone $\widehat{X}$, target sequence $S = (s_1, \ldots, s_L)$, schedule $\kappa$
**Output:** Residue-level scores $\{Q_i\}_{i \in \mathcal{R}}$ and global score $Q$

$\mathcal{R} \leftarrow \textsc{ValidResidues}(\widehat{X}, S)$
$(\widehat{H}_V, \widehat{H}_E) \leftarrow \textsc{BackboneEncoder}(\widehat{X})$
$\widehat{P}_{\mathrm{SS}} \leftarrow \textsc{SecondaryStructureHead}(\widehat{H}_E)$
$C_{\widehat{X}} \leftarrow (\widehat{H}_V, \widehat{H}_E, \widehat{P}_{\mathrm{SS}})$
$\pi \leftarrow \textsc{RandomPermutation}(\mathcal{R})$
$S^{\mathrm{TF}} \leftarrow (N, \ldots, N)$
**for** $m = 1$ to $|\mathcal{R}|$ **do**
  $i \leftarrow \pi_m$
  $t_m \leftarrow \kappa^{-1}\left(\frac{m-1}{|\mathcal{R}|}\right)$
  $E_m \leftarrow \text{RNA-FM}(S^{\mathrm{TF}})$
  $Z_m \leftarrow \textsc{FlowMatchingDecoder}(E_m, C_{\widehat{X}}, t_m)$
  $P_m \leftarrow \mathrm{Softmax}(Z_m)$
  $r_i \leftarrow P_m^{(i)}(s_i)$
  $(S^{\mathrm{TF}})^{(i)} \leftarrow s_i$
**end for**
**for** $i \in \mathcal{R}$ **do**
  $\mathcal{N}_5(i) \leftarrow \{i\} \cup \mathrm{KNN}(i, \widehat{X}_{\mathrm{C4'}}, 4)$
  $Q_i \leftarrow \frac{100}{|\mathcal{N}_5(i)|} \sum_{j \in \mathcal{N}_5(i)} r_j$
**end for**
$Q \leftarrow \frac{1}{|\mathcal{R}|} \sum_{i \in \mathcal{R}} Q_i$
**return** $\{Q_i\}_{i \in \mathcal{R}}, Q$

---

### Supplementary Text

##### Supplementary Text 1. Rosetta-based energetic evaluation of the A29G substitution in RhoBAST

To examine whether the RiboDFM-preferred A29G substitution is energetically compatible with the RhoBAST ligand-binding pocket, we performed Rosetta-based local energetic evaluation of the RhoBAST–MaP555 complex. The experimentally determined RhoBAST-MaP555 structure (PDB ID: 9DXL) was obtained from the Q-BioLiP database[1] and used as the starting structure. The 48-nt RNA chain and the crystallographic coordinates of MaP555 (chemical component ID A1BC9) were extracted from the deposited mmCIF structure.

Because MaP555 is not a standard Rosetta residue, ligand parameters were generated before scoring. The crystallographic ligand coordinates were preserved during ligand preparation and used to initialize both complexes. Aromaticity was assigned using RDKit (v2021.09.3) by partial molecular sanitization while preserving the crystallographic coordinates, after which the ligand was converted to a Rosetta-compatible MOL file with explicitly defined aromatic bonds. Rosetta ligand parameter files were then generated using molfile_to_params.py, with MaP555 represented as residue MP5. The same ligand parameter file and initial crystallographic ligand coordinates were used for all systems.

Wild-type RhoBAST and the A29G single mutant were generated from the same experimental RNA structure using the Rosetta rna_thread application[2], ensuring identical structure-preparation procedures for the two sequences. MaP555 was subsequently placed at its crystallographic position in both complexes. To remove local steric strain introduced by nucleotide substitution while avoiding global rearrangement of the RNA fold, the two complexes were subjected to the same constrained local minimization protocol in Rosetta v2021.16. The RNA backbone was fixed, whereas $\chi$ torsions of residues 16, 29, 31, 33 and 34 and internal ligand torsions were allowed to relax. The rigid-body jump between the RNA and MaP555 was also allowed to minimize. Energy minimization was performed using the lbfgs_armijo_nonmonotone algorithm with a tolerance of 0.001 and a maximum of 500 iterations.

RNA-ligand interaction energetics were evaluated using Rosetta's InterfaceScoreCalculator[3]. The ligand-interface score (interface_delta_X) was calculated as the Rosetta energy difference between the bound RNA–MaP555 complex and the separated RNA and ligand states:

$$\Delta E_{\text{interface}} = E_{\text{bound}} - E_{\text{separated}} \tag{1}$$

where $E_{\text{bound}}$ and $E_{\text{separated}}$ represent the Rosetta scores of the RNA-MaP555 complex and the separated RNA and MaP555 states, respectively. More negative values indicate energetically more favorable predicted RNA-ligand interactions. To quantify the energetic impact of the A29G substitution, we calculated:

$$\Delta\Delta E = \Delta E_{\text{A29G}} - \Delta E_{\text{WT}} \tag{2}$$

All calculations were performed using the Rosetta ligand score function with identical weighting parameters for the wild-type and mutant complexes. Because these scores are reported in Rosetta energy units (REU), they were interpreted as relative interaction scores rather than physical binding free energies in kcal $mol^{-1}$.

After local minimization, the wild-type and A29G complexes yielded interface scores of -19.805 and -19.555 REU, respectively, corresponding to a ΔΔE (A29G - wild type) of +0.250 REU. The close agreement between the two optimized interface scores indicates that the RiboDFM-preferred A29G substitution preserves native-like MaP555-binding energetics within the RhoBAST ligand-binding pocket.

### Supplementary Tables

**Supplementary Table 1. Performance on four RNA design test datasets.** Average values represent the average performance across ten designed sequences per target, whereas top1 values are obtained from the design with the lowest scRMSD among the ten samples, with all metrics reported for the same selected design. Bold and underlined values indicate the best and second-best performance, respectively.

| Evaluation setting | Dataset | Method | Sequence Recovery | sclDDT | scRMSD (Å) | scTM | scINF |
|---|---|---|---|---|---|---|---|
| Average | Mono63 | RiboDFM | <u>0.637</u> | **80.5** | **4.5** | **0.523** | **0.791** |
| | | NA-MPNN | 0.585 | <u>80.2</u> | <u>4.7</u> | <u>0.523</u> | <u>0.786</u> |
| | | gRNAde | 0.551 | 78.0 | 5.4 | 0.502 | 0.772 |
| | | RhoDesign | **0.675** | 55.3 | 14.0 | 0.284 | 0.450 |
| | CASP16 | RiboDFM | **0.549** | **49.3** | **27.5** | **0.286** | **0.543** |
| | | NA-MPNN | <u>0.511</u> | <u>46.6</u> | <u>28.3</u> | <u>0.260</u> | 0.494 |
| | | gRNAde | 0.470 | 46.2 | 28.9 | 0.260 | <u>0.506</u> |
| | | RhoDesign | 0.486 | 30.1 | 36.6 | 0.167 | 0.127 |
| | New38 | RiboDFM | **0.647** | **61.0** | **17.3** | **0.412** | **0.640** |
| | | NA-MPNN | <u>0.582</u> | 55.5 | <u>19.4</u> | 0.349 | <u>0.591</u> |
| | | gRNAde | 0.532 | <u>55.7</u> | 20.2 | <u>0.356</u> | 0.583 |
| | | RhoDesign | 0.563 | 37.9 | 27.2 | 0.214 | 0.251 |
| | Apt35 | RiboDFM | <u>0.667</u> | **71.4** | **5.9** | <u>0.393</u> | **0.683** |
| | | NA-MPNN | **0.691** | <u>71.3</u> | <u>6.0</u> | **0.406** | <u>0.680</u> |
| | | gRNAde | 0.570 | 64.3 | 7.5 | 0.336 | 0.601 |
| | | RhoDesign | 0.627 | 53.3 | 11.8 | 0.272 | 0.383 |
| Top1 | Mono63 | RiboDFM | <u>0.638</u> | <u>83.7</u> | **3.3** | <u>0.566</u> | **0.826** |
| | | NA-MPNN | 0.590 | **83.7** | <u>3.4</u> | **0.580** | <u>0.809</u> |
| | | gRNAde | 0.552 | 81.0 | 4.2 | 0.546 | 0.799 |
| | | RhoDesign | **0.691** | 64.5 | 9.5 | 0.363 | 0.563 |
| | CASP16 | RiboDFM | **0.551** | **52.8** | **22.2** | **0.326** | **0.583** |
| | | NA-MPNN | <u>0.518</u> | <u>50.8</u> | <u>23.6</u> | <u>0.308</u> | 0.530 |
| | | gRNAde | 0.476 | 49.4 | 24.6 | 0.284 | <u>0.548</u> |
| | | RhoDesign | 0.492 | 33.2 | 31.7 | 0.186 | 0.180 |
| | New38 | RiboDFM | **0.646** | **66.4** | **12.9** | **0.467** | **0.701** |
| | | NA-MPNN | <u>0.584</u> | <u>61.2</u> | <u>14.5</u> | <u>0.411</u> | <u>0.636</u> |
| | | gRNAde | 0.527 | 60.9 | 16.0 | 0.402 | 0.627 |
| | | RhoDesign | 0.572 | 42.3 | 22.7 | 0.245 | 0.298 |
| | Apt35 | RiboDFM | <u>0.676</u> | <u>76.3</u> | **4.2** | <u>0.457</u> | <u>0.725</u> |
| | | NA-MPNN | **0.706** | **77.7** | <u>4.2</u> | **0.465** | **0.740** |
| | | gRNAde | 0.572 | 70.5 | 5.4 | 0.389 | 0.672 |
| | | RhoDesign | 0.631 | 57.2 | 9.6 | 0.303 | 0.437 |

**Supplementary Table 2. Comparison of RiboDFM and AlignIF on CASP16 and New38.** Average values represent the average performance across ten designed sequences per target, whereas top1 values are obtained from the design with the lowest scRMSD among the ten samples, with all metrics reported for the same selected design. Bold and underlined values indicate the best and second-best performance, respectively.

| Evaluation setting | Dataset | Method | Sequence Recovery | sclDDT | scTM | scRMSD (Å) | scINF |
|---|---|---|---|---|---|---|---|
| Average | CASP16 | RiboDFM | <u>0.549</u> | <u>49.3</u> | <u>0.286</u> | 27.5 | <u>0.543</u> |
| | | AlignIF (no MStA) | 0.545 | 48.6 | 0.282 | <u>27.0</u> | 0.534 |
| | | AlignIF | **0.550** | **50.0** | **0.296** | **26.6** | **0.544** |
| | New38 | RiboDFM | **0.647** | <u>61.0</u> | <u>0.412</u> | <u>17.3</u> | <u>0.640</u> |
| | | AlignIF (no MStA) | 0.625 | 58.5 | 0.381 | 18.3 | 0.631 |
| | | AlignIF | <u>0.635</u> | **61.4** | **0.422** | **16.6** | **0.654** |
| Top1 | CASP16 | RiboDFM | **0.551** | <u>52.8</u> | **0.326** | **22.2** | <u>0.583</u> |
| | | AlignIF (no MStA) | 0.544 | 51.8 | 0.313 | 22.3 | 0.573 |
| | | AlignIF | <u>0.551</u> | **53.2** | <u>0.324</u> | <u>22.2</u> | **0.598** |
| | New38 | RiboDFM | **0.646** | **66.4** | **0.467** | **12.9** | **0.701** |
| | | AlignIF (no MStA) | 0.628 | 62.2 | 0.423 | 15.0 | 0.660 |
| | | AlignIF | <u>0.640</u> | <u>64.6</u> | <u>0.457</u> | <u>13.8</u> | <u>0.686</u> |

**Supplementary Table 3. Per-sample design results for the RhoBAST aptamer.** Ten sequences generated by each method for RhoBAST (PDB ID: 9DXL) are listed with their sequence recovery and scRMSD to the experimental structure. Structures were predicted independently using trRosettaRNA2[4], AlphaFold 3[5], and Protenix[6]. Red values indicate the RMSD of the structure selected for final comparison, corresponding to the scRMSD-top1 for each design method. For the native sequence, the lowest RMSD among the three structure predictions is highlighted.

| **Method** | **Sample No.** | **Recovery** | **RMSD (Å)** | | |
|---|---|---|---|---|---|
| | | | **trRosettaRNA2** | **AlphaFold 3** | **Protenix** |
| RiboDFM | 1 | 0.833 | 3.8 | 9.6 | 1.4 |
| | 2 | 0.875 | 4.5 | 9.2 | 1.3 |
| | 3 | 0.875 | 2.0 | 9.1 | 1.5 |
| | 4 | 0.875 | 4.5 | 9.2 | 1.3 |
| | 5 | 0.875 | 4.5 | 9.2 | 1.3 |
| | 6 | 0.875 | 4.5 | 9.2 | 1.3 |
| | 7 | 0.875 | 4.5 | 9.2 | 1.3 |
| | 8 | 0.875 | 4.5 | 9.2 | **1.2** |
| | 9 | 0.875 | 4.5 | 9.2 | 1.3 |
| | 10 | 0.875 | 4.5 | 9.2 | 1.3 |
| NA-MPNN | 1 | 0.458 | 18.1 | 19.6 | 18.2 |
| | 2 | 0.583 | 13.8 | 12.2 | 12.4 |
| | 3 | 0.625 | 14.0 | 19.8 | 16.6 |
| | 4 | 0.542 | 11.0 | 20.5 | 8.3 |
| | 5 | 0.563 | 19.9 | 14.2 | 14.4 |
| | 6 | 0.563 | 13.2 | 12.1 | **7.3** |
| | 7 | 0.563 | 24.6 | 19.1 | 8.4 |
| | 8 | 0.625 | 14.3 | 22.1 | 11.0 |
| | 9 | 0.479 | 13.6 | 21.5 | 8.8 |
| | 10 | 0.563 | 17.8 | 18.4 | 18.3 |
| RhoDesign | 1 | 0.688 | 16.8 | 18.1 | 15.6 |
| | 2 | 0.688 | 13.3 | 18.0 | 14.3 |
| | 3 | 0.646 | 16.6 | 18.2 | 14.4 |
| | 4 | 0.688 | 13.3 | 18.0 | 13.9 |
| | 5 | 0.688 | 13.3 | 18.0 | 14.0 |
| | 6 | 0.604 | 17.8 | 19.6 | 17.1 |
| | 7 | 0.625 | 17.9 | 19.8 | 16.9 |
| | 8 | 0.688 | 13.3 | 18.0 | 14.1 |
| | 9 | 0.688 | 12.3 | **11.7** | 13.2 |
| | 10 | 0.667 | 17.5 | 17.6 | 13.9 |
| gRNAde | 1 | 0.500 | **15.3** | 15.9 | 16.3 |
| | 2 | 0.438 | 17.1 | 15.9 | 16.5 |
| | 3 | 0.500 | 18.0 | 15.9 | 15.9 |
| | 4 | 0.500 | 18.0 | 15.9 | 15.9 |
| | 5 | 0.479 | 17.5 | 18.5 | 17.0 |
| | 6 | 0.500 | 18.0 | 15.9 | 15.9 |
| | 7 | 0.458 | 16.2 | 15.8 | 16.1 |
| | 8 | 0.500 | 15.3 | 15.9 | 16.0 |
| | 9 | 0.500 | 15.3 | 15.9 | 16.2 |
| | 10 | 0.458 | 16.2 | 15.7 | 16.1 |
| AlignIF (no MStA) | 1 | 0.583 | 20.7 | 18.7 | 13.7 |
| | 2 | 0.563 | 21.2 | 17.8 | 18.1 |
| | 3 | 0.583 | 20.7 | 18.7 | 13.7 |
| | 4 | 0.583 | 20.7 | 18.7 | **13.5** |
| | 5 | 0.563 | 19.0 | 18.3 | 18.3 |
| | 6 | 0.563 | 20.2 | 18.2 | 17.9 |
| | 7 | 0.583 | 18.6 | 18.9 | 19.0 |
| | 8 | 0.604 | 17.0 | 18.8 | 18.7 |
| | 9 | 0.604 | 18.7 | 18.8 | 18.5 |
| | 10 | 0.563 | 22.8 | 18.3 | 18.7 |
| Native sequence | 1 | 1.000 | 3.6 | 8.4 | **1.6** |

**Supplementary Table 4. RiboDFM ablation performance on four RNA design test datasets.** Average values represent the average performance across ten designed sequences per target, whereas top1 values are obtained from the design with the lowest scRMSD among the ten samples, with all metrics reported for the same selected design. Bold and underlined values indicate the best and second-best performance, respectively.

| Evaluation setting | Dataset | Method | Recovery | sclDDT | scTM | scRMSD (Å) | scINF |
|---|---|---|---|---|---|---|---|
| Average | Mono63 | Baseline | **0.637** | 80.5 | 0.523 | 4.5 | 0.791 |
| | | Autoregressive | 0.565 | 76.0 | 0.464 | 5.7 | 0.748 |
| | | Experimental-only | 0.607 | 79.7 | 0.514 | 4.5 | 0.781 |
| | | w/o SS head | 0.605 | 79.4 | 0.513 | 4.8 | 0.778 |
| | | w/o RNA-FM | 0.628 | **81.4** | **0.541** | **4.4** | **0.794** |
| | CASP16 | Baseline | 0.549 | 49.3 | **0.286** | 27.5 | **0.543** |
| | | Autoregressive | 0.532 | 48.7 | 0.281 | 27.9 | 0.522 |
| | | Experimental-only | 0.550 | 48.4 | 0.273 | 28.0 | 0.523 |
| | | w/o SS head | 0.541 | **49.4** | 0.283 | 27.8 | 0.529 |
| | | w/o RNA-FM | **0.553** | 48.7 | 0.279 | **27.2** | 0.530 |
| | New38 | Baseline | **0.647** | 61.0 | **0.412** | 17.3 | **0.640** |
| | | Autoregressive | 0.603 | 57.4 | 0.370 | 18.7 | 0.611 |
| | | Experimental-only | 0.644 | **61.0** | 0.412 | **17.2** | 0.633 |
| | | w/o SS head | 0.628 | 60.5 | 0.403 | 17.2 | 0.634 |
| | | w/o RNA-FM | 0.647 | 60.3 | 0.397 | 17.2 | 0.629 |
| | Apt35 | Baseline | **0.667** | **71.4** | 0.393 | 5.9 | **0.683** |
| | | Autoregressive | 0.614 | 70.2 | 0.367 | 5.8 | 0.672 |
| | | Experimental-only | 0.634 | 71.0 | 0.388 | 5.9 | 0.674 |
| | | w/o SS head | 0.641 | 70.9 | **0.398** | 6.2 | 0.677 |
| | | w/o RNA-FM | 0.638 | 70.5 | 0.389 | **5.7** | 0.659 |
| Top1 | Mono63 | Baseline | **0.638** | **83.7** | 0.566 | 3.3 | **0.826** |
| | | Autoregressive | 0.569 | 81.7 | 0.541 | 3.6 | 0.788 |
| | | Experimental-only | 0.606 | 83.4 | 0.556 | **3.3** | 0.815 |
| | | w/o SS head | 0.606 | 83.6 | 0.561 | 3.5 | 0.815 |
| | | w/o RNA-FM | 0.630 | 83.6 | **0.570** | 3.5 | 0.811 |
| | CASP16 | Baseline | 0.551 | 52.8 | 0.326 | 22.2 | 0.583 |
| | | Autoregressive | 0.531 | 53.2 | 0.334 | 23.0 | 0.583 |
| | | Experimental-only | 0.545 | 53.0 | 0.325 | 23.2 | 0.567 |
| | | w/o SS head | 0.547 | 52.8 | 0.312 | 23.3 | 0.601 |
| | | w/o RNA-FM | **0.556** | **53.8** | **0.340** | **21.7** | **0.602** |
| | New38 | Baseline | 0.646 | **66.4** | 0.467 | 12.9 | **0.701** |
| | | Autoregressive | 0.606 | 63.3 | 0.436 | 13.5 | 0.663 |
| | | Experimental-only | 0.643 | 65.6 | 0.465 | **12.8** | 0.678 |
| | | w/o SS head | 0.625 | 65.3 | **0.471** | 13.0 | 0.671 |
| | | w/o RNA-FM | **0.649** | 66.3 | 0.464 | 13.0 | 0.684 |
| | Apt35 | Baseline | **0.676** | 76.3 | **0.457** | 4.2 | 0.725 |
| | | Autoregressive | 0.637 | **77.2** | 0.430 | **3.9** | **0.736** |
| | | Experimental-only | 0.641 | 77.1 | 0.441 | 3.9 | 0.720 |
| | | w/o SS head | 0.645 | 75.9 | 0.449 | 4.7 | 0.716 |
| | | w/o RNA-FM | 0.649 | 74.7 | 0.427 | 4.3 | 0.707 |

**Supplementary Table 5. Effect of sampling steps on RiboDFM performance.** Average values represent the average performance across ten designed sequences per target, whereas top1 values are obtained from the design with the lowest scRMSD among the ten samples, with all metrics reported for the same selected design. Bold and underlined values indicate the best and second-best performance, respectively.

| Dataset | Steps | Average | | | | | Top1 | | | | |
|---|---|---|---|---|---|---|---|---|---|---|---|
| | | **Recovery** | **scIDDT** | **scTM** | **scRMSD(Å)** | **scINF** | **Recovery** | **scIDDT** | **scTM** | **scRMSD(Å)** | **scINF** |
| Mono63 | 1 | 0.640 | 77.5 | 0.493 | 5.3 | 0.748 | 0.639 | 81.5 | 0.540 | 3.7 | 0.786 |
| | 2 | 0.639 | 79.1 | 0.509 | 4.9 | 0.764 | 0.640 | 83.5 | 0.561 | 3.4 | 0.816 |
| | 4 | 0.639 | 79.8 | 0.518 | 4.7 | 0.780 | 0.644 | 83.4 | 0.562 | 3.4 | 0.816 |
| | 8 | **0.640** | 80.8 | 0.526 | 4.5 | 0.798 | 0.643 | 83.9 | 0.564 | 3.4 | **0.830** |
| | 16 | 0.638 | 80.5 | 0.522 | 4.4 | 0.796 | 0.644 | 83.6 | 0.563 | 3.4 | 0.826 |
| | 32 | 0.637 | 80.5 | 0.523 | 4.5 | 0.791 | 0.638 | 83.7 | 0.566 | 3.3 | 0.826 |
| | 64 | 0.638 | 80.8 | 0.527 | 4.3 | 0.796 | 0.641 | 83.6 | 0.564 | **3.3** | 0.827 |
| | 128 | 0.639 | 80.9 | 0.526 | **4.3** | 0.798 | 0.643 | 83.7 | 0.557 | 3.3 | 0.826 |
| | 256 | 0.638 | 80.8 | 0.525 | 4.4 | 0.797 | 0.644 | 83.7 | 0.562 | 3.4 | 0.827 |
| | 512 | 0.639 | 80.9 | **0.528** | 4.3 | **0.803** | 0.641 | 83.5 | **0.567** | 3.3 | 0.827 |
| | 1024 | 0.638 | **80.9** | 0.525 | 4.3 | 0.800 | **0.645** | **83.9** | 0.561 | 3.3 | 0.825 |
| CASP16 | 1 | **0.556** | 44.6 | 0.257 | 30.1 | 0.420 | **0.559** | 48.0 | 0.282 | 25.9 | 0.472 |
| | 2 | 0.552 | 46.8 | 0.267 | 29.1 | 0.473 | 0.553 | 51.3 | 0.315 | 23.5 | 0.552 |
| | 4 | 0.552 | 47.9 | 0.275 | 28.6 | 0.507 | 0.554 | 51.6 | 0.308 | 24.0 | 0.567 |
| | 8 | 0.549 | 48.7 | 0.285 | 27.9 | 0.526 | 0.554 | 52.6 | 0.328 | 23.4 | 0.587 |
| | 16 | 0.548 | 49.0 | 0.284 | 28.1 | 0.533 | 0.549 | 51.2 | 0.313 | 24.0 | 0.569 |
| | 32 | 0.549 | 49.3 | 0.286 | 27.5 | 0.543 | 0.551 | 52.8 | 0.326 | 22.2 | 0.583 |
| | 64 | 0.548 | 49.1 | 0.284 | **27.4** | 0.539 | 0.551 | 52.7 | 0.330 | **21.8** | 0.605 |
| | 128 | 0.547 | 49.3 | 0.287 | 27.5 | 0.539 | 0.549 | 53.1 | 0.310 | 22.2 | 0.593 |
| | 256 | 0.548 | 49.2 | 0.286 | 27.6 | 0.542 | 0.544 | 52.9 | 0.328 | 22.3 | 0.583 |
| | 512 | 0.548 | **49.8** | **0.290** | 27.8 | **0.557** | 0.549 | **53.6** | **0.339** | 23.0 | **0.620** |
| | 1024 | 0.546 | 49.3 | 0.283 | 27.7 | 0.550 | 0.552 | 52.9 | 0.324 | 21.8 | 0.608 |
| New38 | 1 | 0.645 | 56.3 | 0.374 | 19.6 | 0.553 | 0.643 | 61.6 | 0.423 | 15.1 | 0.626 |
| | 2 | 0.645 | 58.3 | 0.390 | 18.6 | 0.584 | 0.652 | 65.3 | 0.467 | 14.2 | 0.682 |
| | 4 | 0.647 | 59.4 | 0.400 | 18.0 | 0.617 | 0.651 | 65.6 | 0.467 | 13.5 | 0.682 |
| | 8 | 0.646 | 60.9 | 0.413 | 17.3 | 0.637 | 0.649 | 66.5 | 0.474 | 13.1 | 0.684 |
| | 16 | 0.646 | 61.1 | 0.412 | 17.2 | 0.643 | 0.652 | 66.8 | 0.474 | **12.4** | 0.703 |
| | 32 | 0.647 | 61.0 | 0.412 | 17.3 | 0.640 | 0.646 | 66.4 | 0.467 | 12.9 | 0.701 |
| | 64 | 0.647 | 60.9 | 0.410 | 17.3 | 0.641 | 0.648 | **67.6** | **0.487** | 12.6 | **0.710** |
| | 128 | 0.645 | 61.5 | 0.418 | 17.0 | 0.647 | **0.654** | 66.5 | 0.475 | 12.9 | 0.700 |
| | 256 | 0.645 | 61.1 | 0.417 | 17.2 | 0.646 | 0.644 | 66.3 | 0.470 | 13.0 | 0.697 |
| | 512 | **0.647** | **61.9** | **0.423** | 16.9 | **0.654** | 0.650 | 65.9 | 0.479 | 12.7 | 0.690 |
| | 1024 | 0.647 | 61.8 | 0.422 | **16.8** | 0.650 | 0.646 | 66.8 | 0.473 | 12.5 | 0.699 |
| Apt35 | 1 | 0.664 | 69.2 | 0.368 | 6.7 | 0.650 | 0.666 | 73.2 | 0.413 | 4.8 | 0.699 |
| | 2 | 0.663 | 70.5 | 0.383 | 6.1 | 0.657 | 0.671 | 76.6 | 0.447 | 4.0 | 0.730 |
| | 4 | 0.665 | 71.2 | 0.391 | 6.1 | 0.670 | **0.680** | 76.4 | 0.452 | 4.0 | 0.711 |
| | 8 | 0.664 | 71.1 | 0.388 | 6.0 | 0.676 | 0.665 | 76.1 | 0.445 | 4.1 | 0.732 |
| | 16 | 0.660 | 71.3 | 0.392 | 5.9 | 0.681 | 0.667 | 75.7 | 0.440 | 4.4 | 0.731 |
| | 32 | **0.667** | 71.4 | 0.393 | 5.9 | 0.683 | 0.676 | 76.3 | **0.457** | 4.2 | 0.725 |
| | 64 | 0.664 | 71.4 | 0.396 | 6.0 | 0.684 | 0.662 | **76.8** | 0.447 | **4.0** | **0.734** |
| | 128 | 0.663 | 71.5 | 0.392 | 6.0 | 0.686 | 0.676 | 76.1 | 0.437 | 4.4 | 0.711 |
| | 256 | 0.664 | **71.9** | **0.399** | **5.7** | **0.688** | 0.675 | 76.2 | 0.440 | 4.1 | 0.725 |
| | 512 | 0.664 | 71.6 | 0.399 | 5.8 | 0.685 | 0.668 | 75.3 | 0.433 | 4.3 | 0.708 |
| | 1024 | 0.661 | 71.3 | 0.395 | 6.1 | 0.686 | 0.672 | 76.6 | 0.449 | 4.1 | 0.729 |

### Supplementary Figures

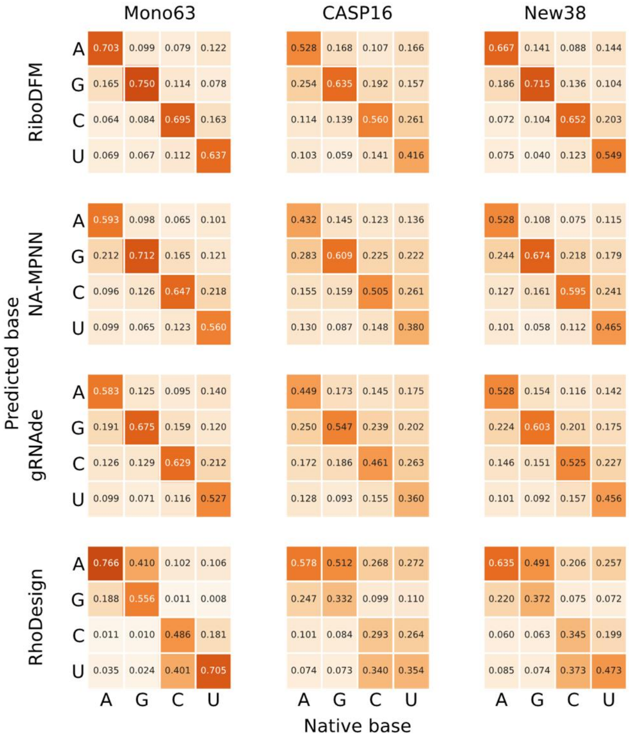


**Supplementary Figure 1. Nucleotide prediction confusion matrices across the three test datasets.** Each matrix summarizes the nucleotide prediction patterns for a given method and dataset. Columns correspond to native nucleotide identities, whereas rows correspond to predicted nucleotide identities. Each cell reports the fraction of nucleotides of the native type indicated by the column that are predicted as the nucleotide type indicated by the row. Values are normalized within each native nucleotide class such that each column sums to one.

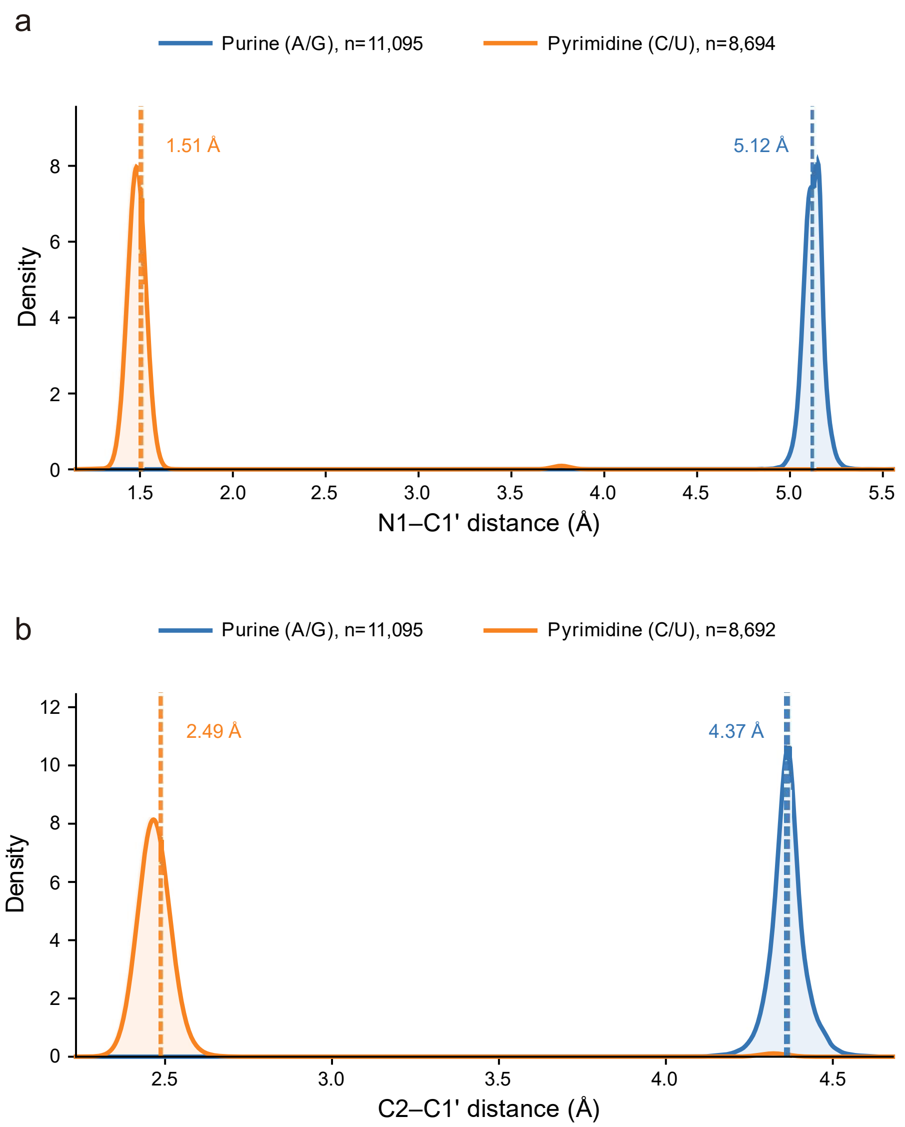


**Supplementary Figure 2. Distributions of base–sugar distances for purines and pyrimidines across the three test datasets.** Distributions of (a) N1-C1′ and (b) C2-C1′ distances for purine (A/G) and pyrimidine (C/U) nucleotides pooled across the three benchmark datasets. Dashed lines indicate the mean distances of the pooled distributions.

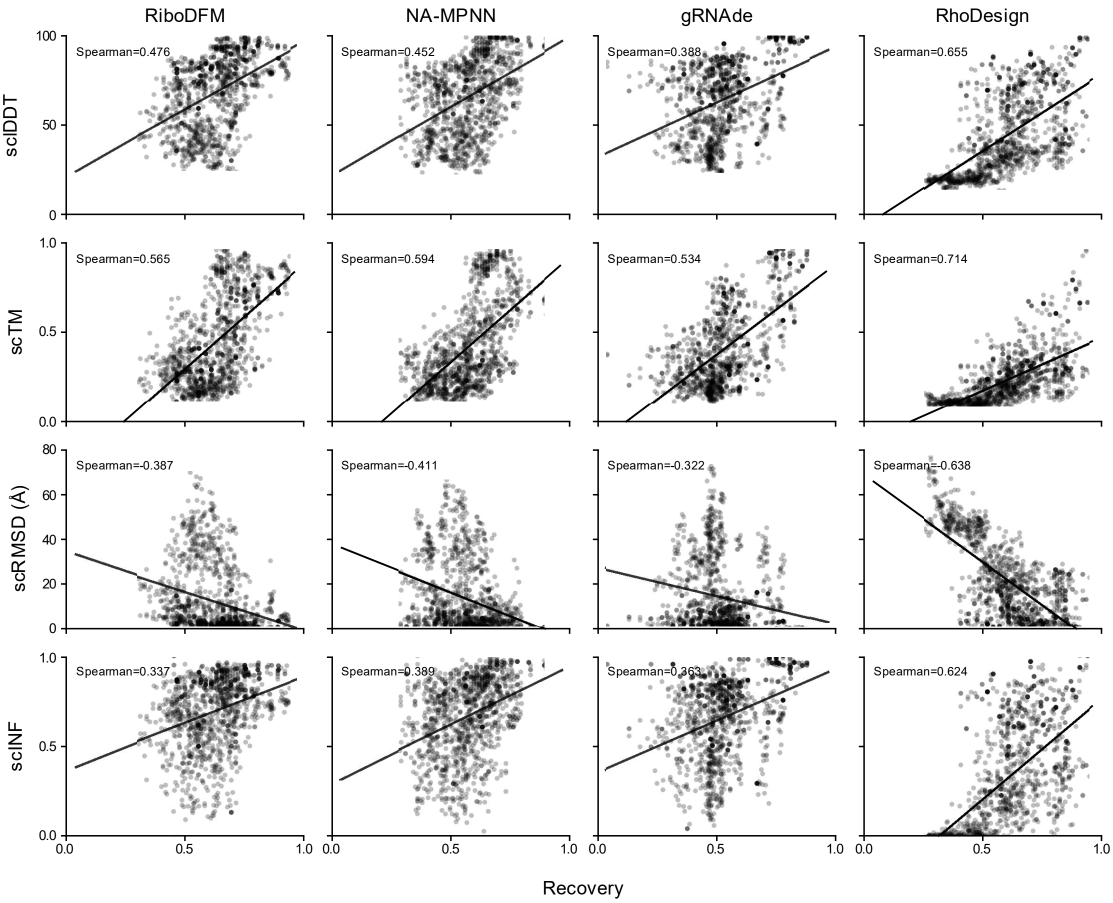


**Supplementary Figure 3. Relationship between sequence recovery and structural self-consistency on all three datasets.**

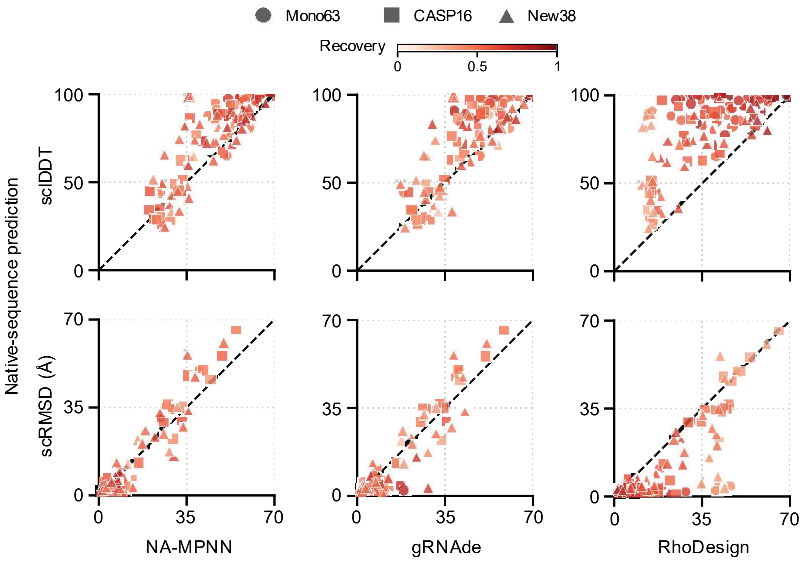


**Supplementary Figure 4. Top1 NA-MPNN, gRNAde and RhoDesign designs (lowest scRMSD) compared with native-sequence predictions.** Colors indicate sequence recovery and shapes denote datasets.

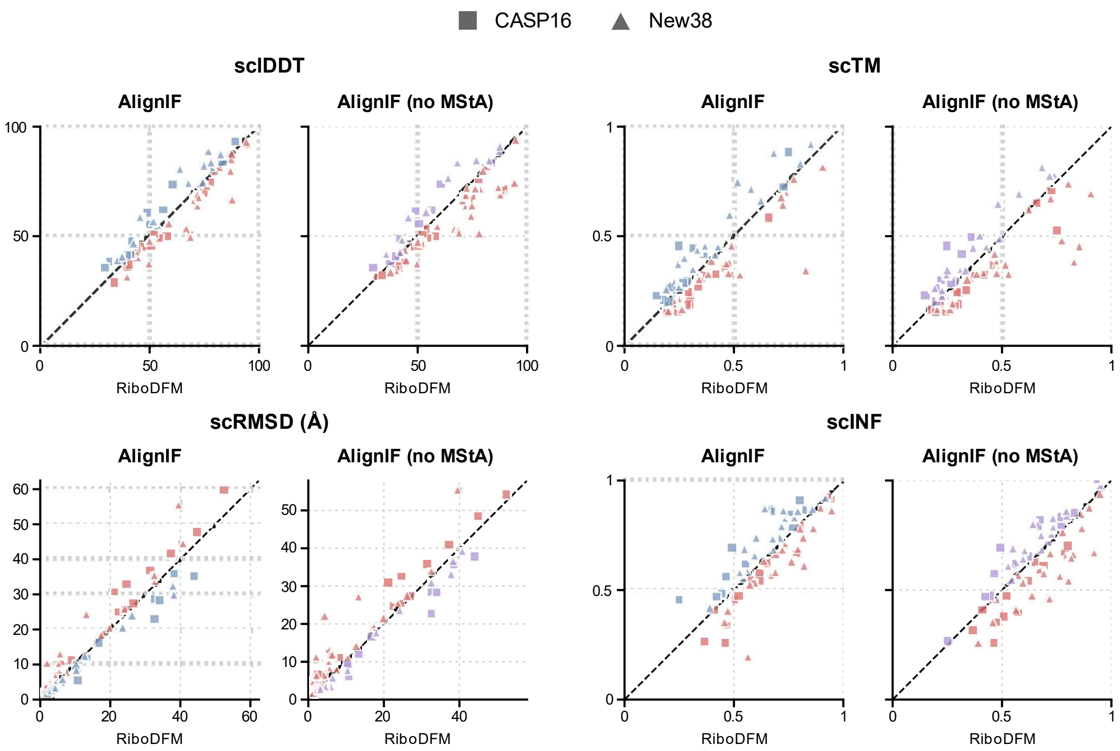


**Supplementary Figure 5. Target-wise Top1 (lowest scRMSD) comparison between RiboDFM and AlignIF in terms of 3D structural self-consistency.**

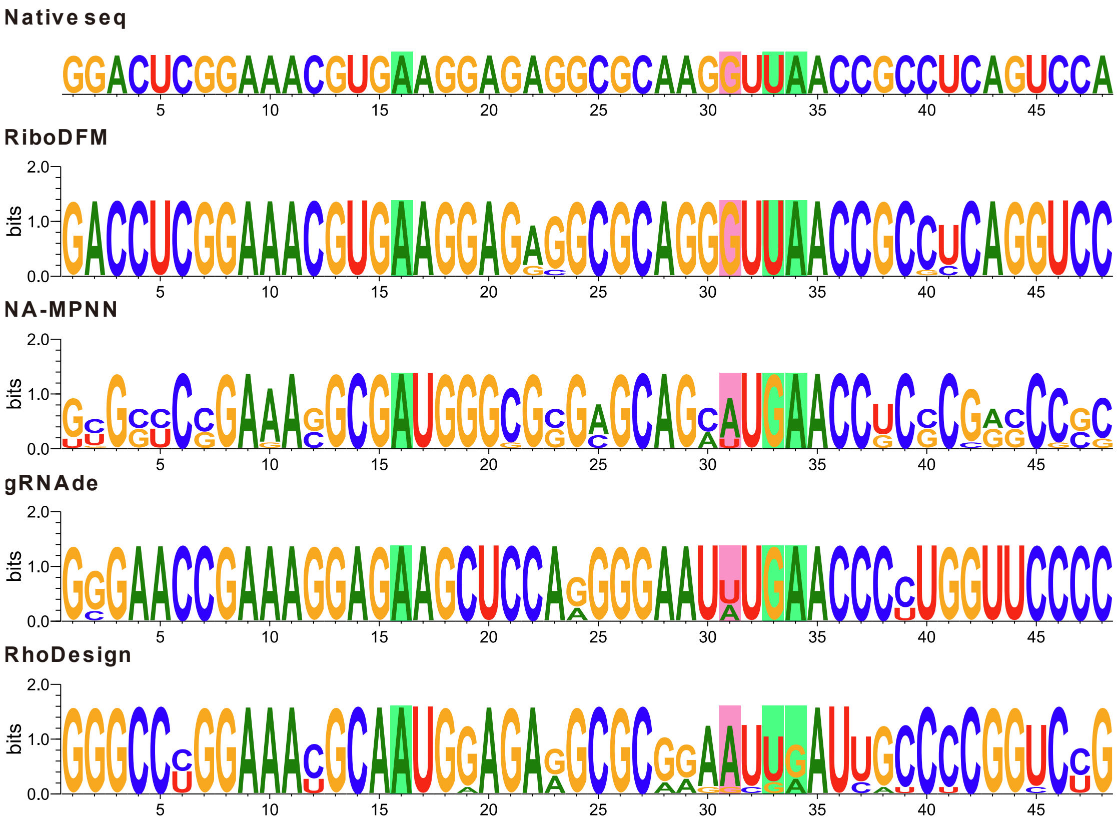


**Supplementary Figure 6. Sequence conservation of designs for the RhoBAST aptamer.** The native sequence of RhoBAST (PDB ID: 9DXL) is shown above sequence logos constructed from ten sequences generated by RiboDFM, NA-MPNN, gRNAde and RhoDesign. Letter heights indicate nucleotide frequencies at individual positions. Positions 16, 31, 33 and 34 are highlighted with distinct background colors.

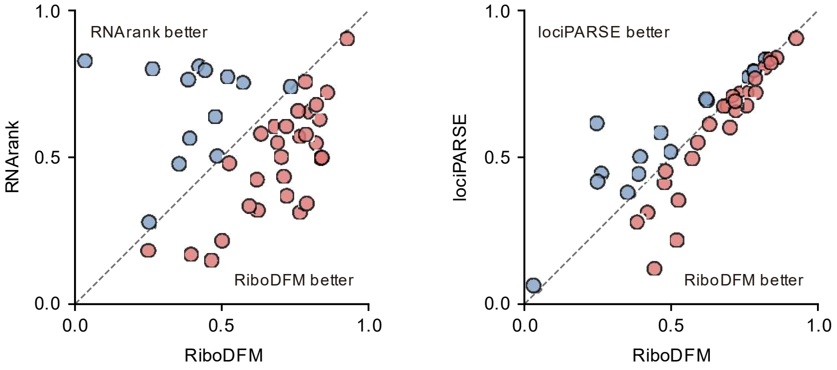


**Supplementary Figure 7. Target-wise global Spearman comparison of zero-shot RNA structure quality assessment by RiboDFM with RNArank and lociPARSE.**

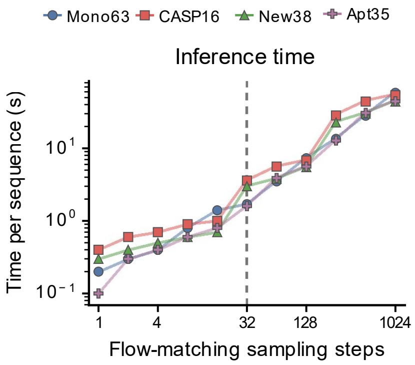


**Supplementary Figure 8. Effects of sampling steps on inference time across four test sets.**

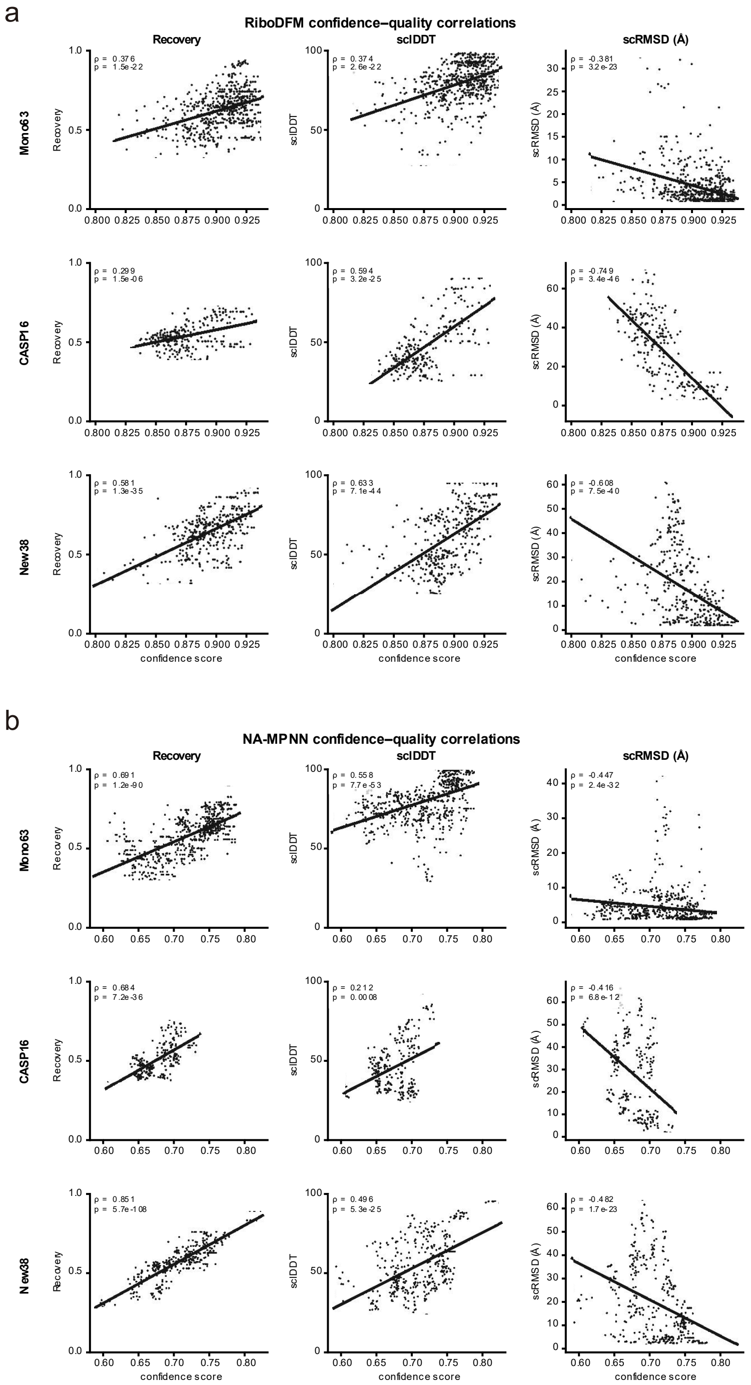


**Supplementary Figure 9. Correlations between confidence scores and design quality.** a,b, Scatter plots of confidence scores against sequence recovery, sclDDT and scRMSD for RiboDFM (a) and NA-MPNN (b) on Mono63, CASP16 and New38.
